# *De Novo* Discovery of High Affinity Peptide Binders for the SARS-CoV-2 Spike Protein

**DOI:** 10.1101/2020.09.29.317131

**Authors:** Sebastian Pomplun, Muhammad Jbara, Anthony J. Quartararo, Genwei Zhang, Joseph S. Brown, Yen-Chun Lee, Xiyun Ye, Stephanie Hanna, Bradley L. Pentelute

## Abstract

The β-coronavirus SARS-CoV-2 has caused a global pandemic. Affinity reagents targeting the SARS-CoV-2 spike protein, the most exposed surface structure of the virus, are of interest for the development of therapeutics and diagnostics. We used affinity selection-mass spectrometry for the rapid discovery of synthetic high affinity peptide binders for the receptor binding domain (RBD) of the SARS-CoV-2 spike protein. From library screening with 800 million synthetic peptides, we identified three sequences with nanomolar affinities (dissociation constants *K*_d_ = 80 to 970 nM) for RBD and selectivity over human serum proteins. Picomolar RBD concentrations in biological matrix could be detected using the biotinylated lead peptide in ELISA format. These peptides might associate with the SARS-CoV-2-spike-RBD at a site unrelated to ACE2 binding, making them potential orthogonal reagents for sandwich immunoassays. We envision our discovery as a robust starting point for the development of SARS-CoV-2 diagnostics or conjugates for virus directed delivery of therapeutics.

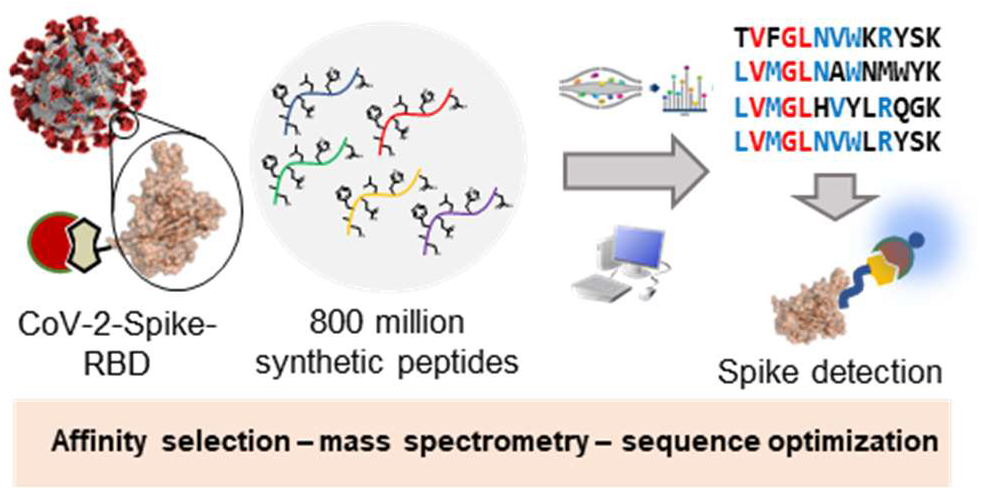

## Introduction

Since the end of 2019, the severe acute respiratory syndrome coronavirus 2 (SARS-CoV-2) has caused the global coronavirus disease 2019 (COVID-19) pandemic. With 30 million cases and over 950,000 deaths, SARS-CoV-2 has spread further than SARS-CoV-1 or middle east respiratory syndrome (MERS).^1,2^ There is a crucial need for diagnostic testing of SARS-CoV-2 to improve outbreak containment.^3,4^ Currently, the reverse-transcriptase polymerase chain reaction (RT-PCR) method is the gold standard of SARS-CoV-2 detection.^5,6^ Serologic detection of patient derived antibodies can be used to track SARS-CoV-2 progression and immunity, but has limited early detection ability.^6–8^ Direct detection of SARS-CoV-2 has been proposed in scalable, rapidly deployed formats, but often suffers from low sensitivity that limits effectivity in general population testing.^9,10^ Thus, the discovery of additional reagents to enable early and rapid SARS-CoV-2 detection and/or neutralization is critical.

Recognition of viral surface proteins by high affinity reagents represents a promising strategy for virus detection or neutralization. Coronaviruses display multiple copies of spike protein on their surface as the most exposed moiety of the virus.^11,12^ SARS-CoV-2 (as well as SARS-CoV-1) binds with high affinity to human angiotensin converting enzyme 2 (ACE2) receptors as a primary mechanism for initiating cell invasion.^13,14^ Several proteins that target SARS-CoV-2 spike have been described. Soluble human and modified ACE2 show high affinity to the SARS-CoV-2 spike protein receptor binding domain (RBD) and neutralizing activity in live virus infection models.^15,16^ Computational design led to the discovery of high affinity miniprotein binders for RBD.^17^ Also, several neutralizing antibodies and nanobodies binding to the SARS-CoV-2-spike-RBD have been described.^18,19^

Peptide sequences with high affinity and selectivity for the SARS-CoV-2 spike protein and its RBD could be advanced for the development of new diagnostic or therapeutic modalities. Peptides have been previously investigated as potential SARS-CoV-2 antiviral agents. For example, long α-helical peptides binding to the S2 unit of the coronavirus spike protein have been described as potent fusion inhibitors.^20^ Computational and experimental studies supported targeting the spike RBD with linear peptides derived from the ACE2 N-terminus.^21,22^ Investigations in our laboratory, however, indicated that peptides 12 to 23 residues long derived from the ACE2 N-terminus do not associate with high affinity to SARS-CoV-2-spike-RBD expressed in human cells.^23^ With straightforward handling, preparation, and late-stage modification,^24–26^ peptides are attractive potential candidates for point-of-care diagnostics.

Here, we report the discovery of synthetic peptides (13 residues) with nanomolar affinity for the SARS-CoV-2-spike-RBD. We leveraged a combinatorial affinity selection-mass spectrometry (AS-MS) platform^27^ for the rapid identification of sequences that display high affinity toward RBD with selectivity over human proteins. After synthesis of the identified peptides, validation of their selective binding activity was observed using biolayer interferometry (BLI) and a magnetic bead pulldown assay. In an enzyme-linked immunosorbent assay (ELISA), these peptides detected picomolar concentrations of RBD mixed in a complex biological matrix.

## Results and discussion

Peptides with a shared sequence motif were enriched and identified by affinity selection-mass spectrometry. SARS-CoV-2-spike-RBD was biotinylated and immobilized on streptavidin-coated magnetic beads. Following an AS-MS protocol recently established by our group,^27^ we screened four peptide libraries with ~200 million members each against the immobilized spike RBD. The peptide library design was X_12_K, where X = any canonical amino acid except Ile and Cys. In parallel, with the same library we performed an enrichment with the anti-hemagglutinin monoclonal antibody 12ca5 to identify non-specific binders, which appeared in both selections. Unbound peptides were removed by three washes with 1x PBS; potential binders were eluted with the denaturant guanidinium hydrochloride and analyzed by nano liquid chromatography-tandem mass spectrometry (nanoLC-MS/MS). The resultant MS spectra were visualized with the sequencing software PEAKS 8.5 and further refined with a Python script that nominated variants matching the library design.^28^

From the ~800 million screened peptides, three peptide sequences were identified (**1-3**, Figure 1a). An extracted ion chromatogram was created for each peptide, revealing each was selectively enriched against the SARS-CoV-2-spike-RBD and not the 12ca5 off-target control (SI Figure 2-4). The three identified peptides shared a common motif at the N-terminus: *V*GL (red letters in Figure 1A). For the additional six positions, we observed enrichment for specific residues as indicated by blue letters (Figure 1A). Based on the similarity of the three sequences, the residues with the highest positional frequencies were combined into a consensus sequence (**4**, Figure 1a).

**Figure 1.**
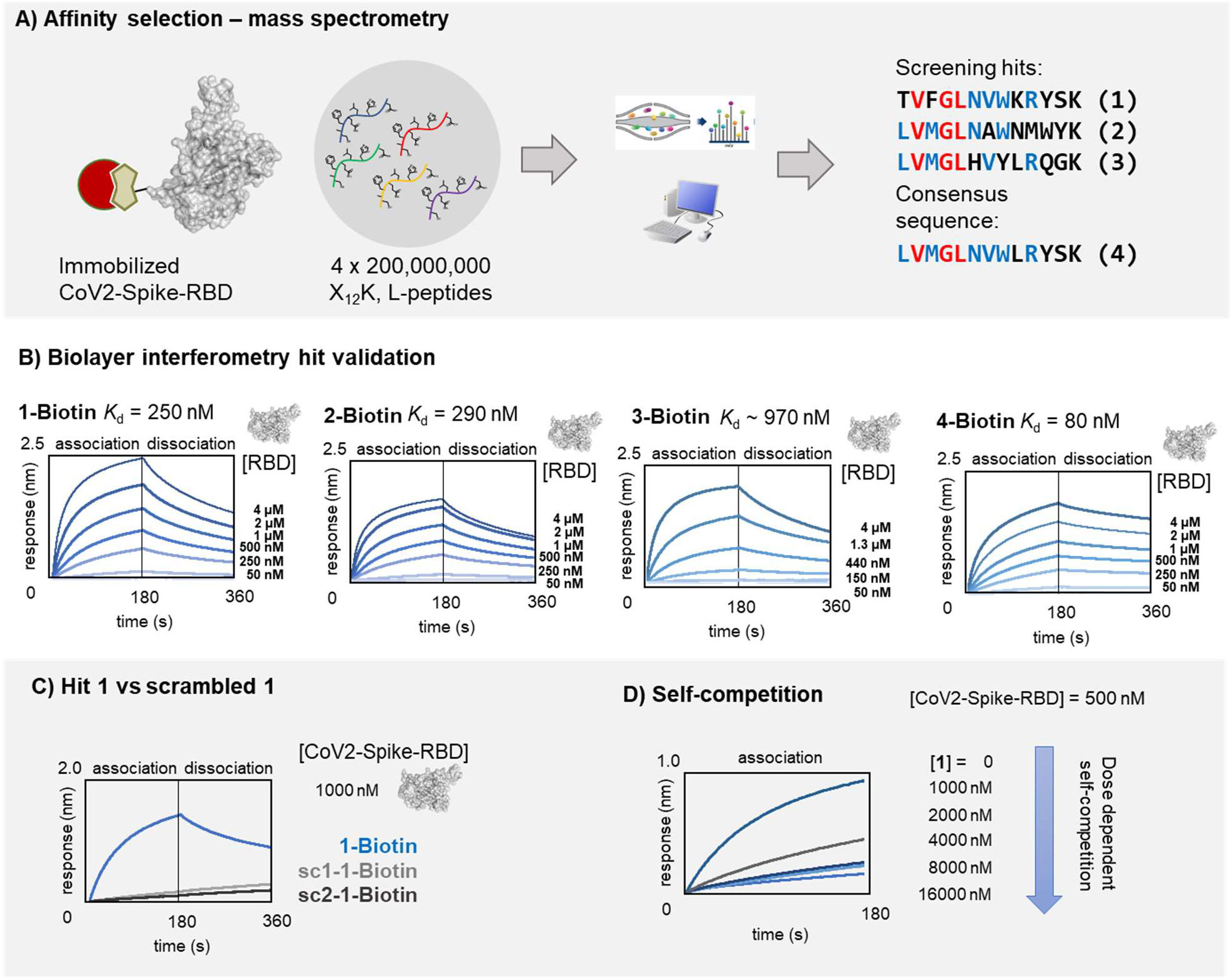
SARS-CoV-2-spike-RBD binding peptides with nanomolar affinity were identified by affinity selection-mass spectrometry. A) Schematic representation of the AS-MS workflow and enriched sequences. In brief: biotinylated SARS-CoV-2-spike-RBD was immobilized on magnetic streptavidin beads and then incubated with peptide libraries. Unbound members were removed by washing. Peptides bound to SARS-CoV-2-spike-RBD were eluted and analyzed by nanoLC-MS/MS. B) BLI curves for association/dissociation of **1, 2,** and **4** to SARS-CoV-2-spike-RBD (in kinetic buffer: 1x PBS, pH = 7.2, 0.1% bovine serum albumin, 0.02% Tween-20). While peptide **4** had somewhat higher affinity, peptide **1**, compared to **2** and **4,** had the best solubility and was used for all further investigations. Peptides **2** and **4** precipitated within hours at concentrations greater than 10 μM. Kinetic binding results are reported in SI Table 1. C) BLI curves for **1** (blue line) and scrambled analogs of **1** (light and dark grey lines respectively, sc1: GSVKRWLTYVKNFK and sc2: RFYVTKGWSNKVLK). D) Self-competition analysis (BLI association) of **1** to SARS-CoV-2-spike-RBD: **1-biotin** immobilized on BLI tips was dipped into solutions containing SARS-CoV-2-spike-RBD and **1** ([RBD] = 500 nM; [**1**] = 0 – 16 μM). Increasing the concentration of **1** in solution causes less free RBD available in solution (due to RBD-**1** complex formation) and results in a concentrationdependent decrease in BLI response.

The identified peptide sequences bound SARS-CoV-2-spike-RBD with nanomolar affinity. To validate the identified sequences, we synthesized biotinylated peptides **1, 2, 3** and **4**. The compounds were immobilized on streptavidin-coated biolayer interferometry (BLI) tips and used to measure association and dissociation of SARS-CoV-2-spike-RBD at different concentrations. Peptides **1, 2 and 3** bound the SARS-CoV-2-spike-RBD with apparent dissociation constants, *K*_d_, of 250 nM, 290 nM, and 900 nM respectively (Figure 1B). The consensus peptide **4** bound the SARS-CoV-2-spike-RBD with *K*_d_ = 80 nM, a ~3-fold improvement over the originally identified hit peptides (Figure 1B). To determine whether the binding is sequence-specific, two scrambled variants of **1** (sc1 and sc2) were prepared and tested by BLI, and no association was observed in either case (Figure 1C). We also tested the association to 12ca5: compared to a known positive control 12ca5 binder, peptide **1** showed minimal association (SI Figure 5). While **4** had the highest affinity to SARS-CoV-2-spike-RBD, peptide **1** displayed the highest solubility of the tested peptides and was used for all further investigations.

Peptide **1** associates to a specific site on the SARS-CoV-2-spike-RBD as determined by a self-competition binding assay. We immobilized **1-biotin** on BLI tips and dipped them into solutions containing SARS-CoV-2-spike-RBD and **1** ([RBD] = 500 nM; [**1**] = 0 – 16 μM). With an increasing concentration of **1** in solution, we observed a concentration-dependent decrease in binding. With less free RBD available in solution, less RBD associates to the BLI tip coated in **1** (Figure 1C).

Peptide **1** does not interfere with ACE2 binding to the RBD. For this experiment, we immobilized ACE2 on BLI tips and dipped them into solutions of SARS-CoV-2-spike-RBD and **1** ([RBD] = 100 nM; [**1**] = 0 – 62 μM, SI Figure 6). We did not observe a decrease in binding response. This can be explained by peptide **1** not having sufficient affinity to block binding of RBD to ACE2 or that peptide **1** might bind to a site on the RBD separate from ACE2. In the latter case, ACE2 and **1-biotin** might form an orthogonal ligand pair useful for spike-RBD sandwich immunoassays.

Alanine scanning mutagenesis of sequence **1** was used to determine the importance of the common motif (*V*GL) and other residues in binding to RBD (Figure 2). Mutations V2A and G4A ablated binding to SARS-CoV-2-spike-RBD, confirming their contributions to the motif identified (Figure 1A). Mutations of hydrophobic residues reduced the peptide binding affinity. Specifically, the mutants F3A, L5A, and W8A decreased the binding by 5 to 13-fold. Position 3 in the other identified peptides contained a methionine suggesting a hydrophobic residue driving is important at this site. Similarly, position 8 needs to be either tryptophan or tyrosine, indicating that an aromatic group drives a higher affinity interaction.

**Figure 2.**
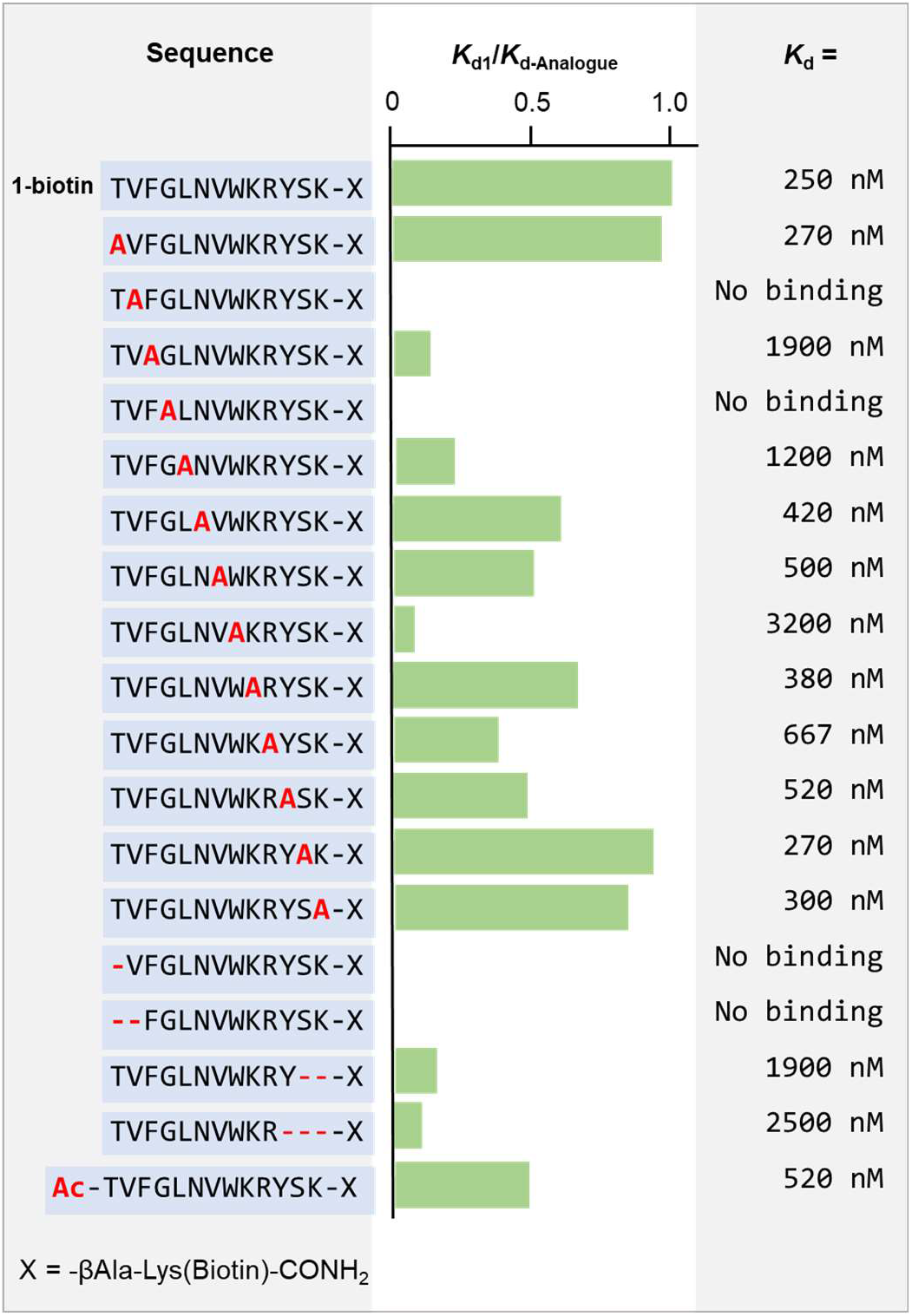
Alanine scanning and sequence truncations of 1 reveal binding hotspots. Binding to SARS-CoV-2-spike-RBD of alanine mutants and truncated peptides was measured by BLI as detailed in Figure 1. Ratios between binding of original sequence **1** and each mutant, respectively, are shown as green bars. Individual steady state *K*_d_ values are shown in the right column. Kinetic binding results are reported in SI Table 1.

To assess the importance of sequence length, we synthesized N-terminal and C-terminal truncations of **1**. A one residue truncation from the N-terminus abolished binding, confirming the frame dependency of the binding motif. N-terminal acetylation reduced the binding affinity to *K*_d_ = 520 nM, indicating the N-terminal amino group is important. Truncations from the C-terminus decreased binding ~10-fold.

Peptide **1** binds to the spike proteins of SARS-CoV-1 and MERS coronaviruses. Patient-derived antibodies often bind SARS-CoV-1 and −2 selectively, with less binding activity against MERS.^29^ However, the spike protein shares some degree of sequence and structural similarity among several species of the β-coronavirus family.^30^ We tested the binding of peptide **1** to the RBDs of SARS-CoV-1 and MERS, and observed *K*_d_ of 600 nM and 120 nM, respectively (Figure 3A). The *K*_d_ values were found in the same order of magnitude as the binding of **1** to SARS-CoV-2-spike-RBD.

**Figure 3.**
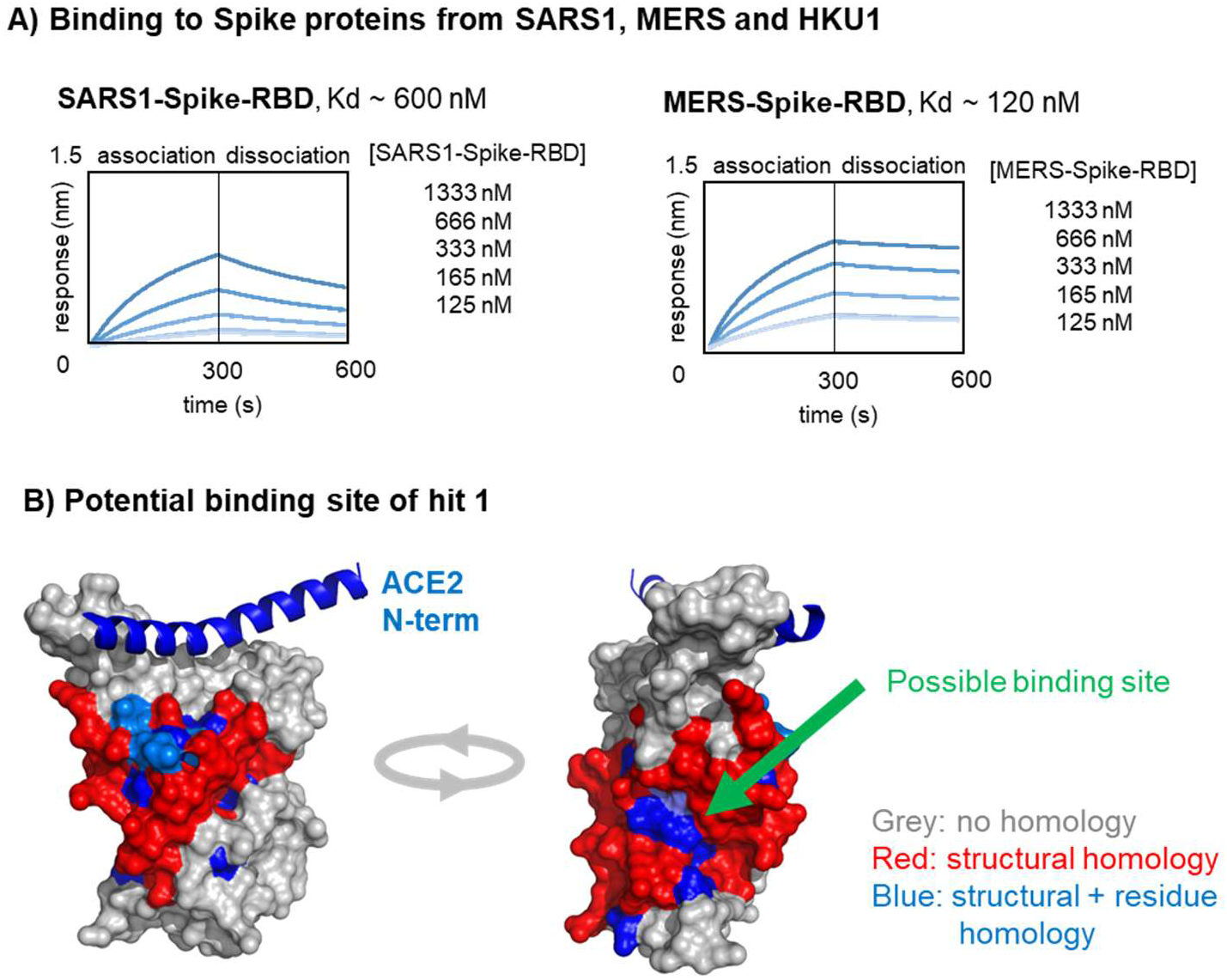
Peptide 1 binds to SARS-CoV-1-Spike RBD and MERS-Spike-RBD. A) The binding of **1-Biotin** to SARS-CoV-1-Spike-RBD and MERS-RBD,was determined by BLI. B) A structural overlay of SARS-CoV-2-spike-RBD and MERS-spike-RBD was performed with the software PyMol (using PDB structures 6vw1 and 6c6z). Regions with homology of secondary structure between the two proteins are colored in red. The homologous regions were manually analyzed to identify positions with identical residues in both proteins: blue. Since **1-biotin** binds to both with comparable affinity, the binding site could potentially be in a region with high homology.

We also investigated the binding of **1** against the spike protein of HKU1, which is an endemic human coronavirus.^29,31^ Since the receptor-binding domain of this protein was not commercially available, we compared the binding of **1** to the HKU1 spike S1 protein subunit and to the SARS-CoV-2 spike S1 protein subunit (both expressed in mammalian cells). The binding of **1** to SARS-CoV-2-S1 demonstrated a *K*_d_ of 40 nM, but showed considerably weaker association to the HKU1-S1 protein (SI Figure 7). The stronger binding affinity of **1** to SARS-CoV-2-S1 (40 nM) compared to SARS-CoV-2-spike-RBD (250 nM) could arise from reduced conformational freedom of the RBD as a part of the S1 spike protein compared to the RBD alone. Alternatively, the difference could be explained by potential differences in quality of recombinant expression of the different commercial proteins.

Significant binding was observed with MERS and **1**, suggesting a similar binding site might be recognized. We were intrigued by the retained binding affinity of **1** to MERS as SARS-CoV-1, SARS-CoV-2, and MERS RBDs have a high level of structural homology. On a residue level, however, only SARS-CoV-1 and SARS-CoV-2 share a high level of sequence homology (~74%) while MERS and SARS-CoV-2-spike-RBD s share only ~24%. Analyzing the overlay of these proteins via PyMOL, we found only two patches with residue homology large enough to associate with a peptide binder (colored in blue in Figure 3B). One of these patches is located in proximity to the ACE2 binding site, the other on the opposite side of the RBD. Since no perturbation of the ACE2 binding to RBD was observed in the presence of **1**, the patch on the opposite side of the RBD is suggested as a possible binding site for **1**.

SARS-CoV-2-spike-RBD can be selectively enriched from human serum proteins using immobilized peptide **1**. To investigate the binding specificity of **1** to RBD in a biological matrix, we added SARS-CoV-2-spike RBD to human serum (10% in 1xPBS) and incubated with magnetic beads displaying **1-biotin** (Figure 4A). We removed the supernatant, washed the beads, and treated with urea to elute bound proteins. We analyzed the urea fraction by SDS-PAGE and observed selective enrichment of SARS-CoV-2-spike-RBD; no other proteins were detected (Figure 4B). The selective binding of **1** to SARS-CoV-2-spike-RBD and isolation from complex biological media is a promising feature for potential diagnostic applications or the use of this peptide as a delivery agent for antiviral payloads.

**Figure 4.**
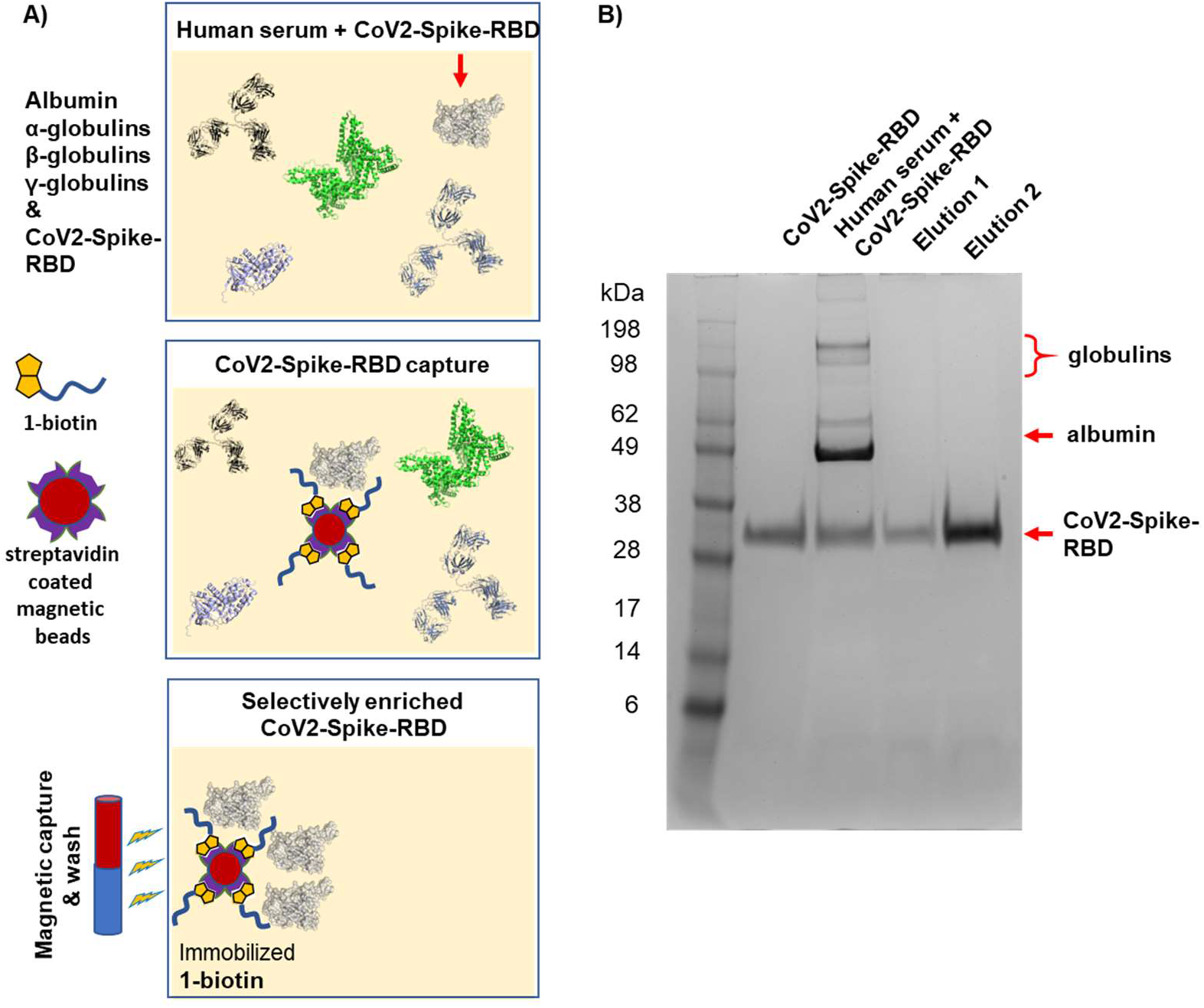
CoV2-Spike-RBD can be selectively enriched from human serum proteins. A) Schematic representation of the pulldown of SARS-CoV-2-spike-RBD from human serum: spike RBD was added to human serum (RBD: 0.27 mg/mL, 10% human serum, 1x PBS), and the mix was incubated with magnetic beads (MyOne Dynabeads) displaying **1-biotin** (1 h, 4 °C). The supernatant (containing nonbinding proteins) was removed, and the beads washed with 1x PBS (3 x 1 mL). Bound proteins were eluted with 6 M urea (elution 1: 50 μL, 30 sec, elution 2: 50 μL, 120 sec) and analyzed by SDS PAGE (B). The gel shows (from left to right): 1) molecular weight ladder; 2) purified SARS-CoV-2-spike-RBD (1 μg); 3) Human serum mixed with SARS-CoV-2-spike RBD; 4) Elution 1 (30 μL of elution sample 1); 5) elution 2 (30 μL of elution sample 2). The analysis was performed using BoltTM 4-12% Bis-Tris Plus Gels (10-wells), 165 V for 36 min, utilizing pre-stained Invitrogen SeeBlueTM Plus2 molecular weight standard with BoltTM LDS Sample Buffer (4X).

Picomolar concentrations of RBD in biological matrix were detected by **1-biotin** in an enzyme-linked immunosorbent assay (ELISA) format. We performed an ELISA detection by immobilizing different concentrations of RBD (100 nM – 100 fM) onto the ELISA plate in fetal bovine serum (FBS). We then incubated the plate with **1-biotin** (100 nM), followed by streptavidin-HRP (horseradish peroxidase) and tetramethylbenzidine (TMB) substrate (Figure 5A). We detected SARS-CoV-2-spike-RBD at 100 nM and 100 pM concentrations with a signal significantly stronger than the background (Figure 5B). The background signal caused by the peptide in absence of RBD on the plate could be caused by a certain degree of unspecific binding of **1-biotin** to components of the biological matrix. Initial experiments with SARS-CoV-2-spike-RBD dissolved in human serum also resulted in a dose response detection with a significant signal at nanomolar RBD concentration (SI Figure 10).

**Figure 5.**
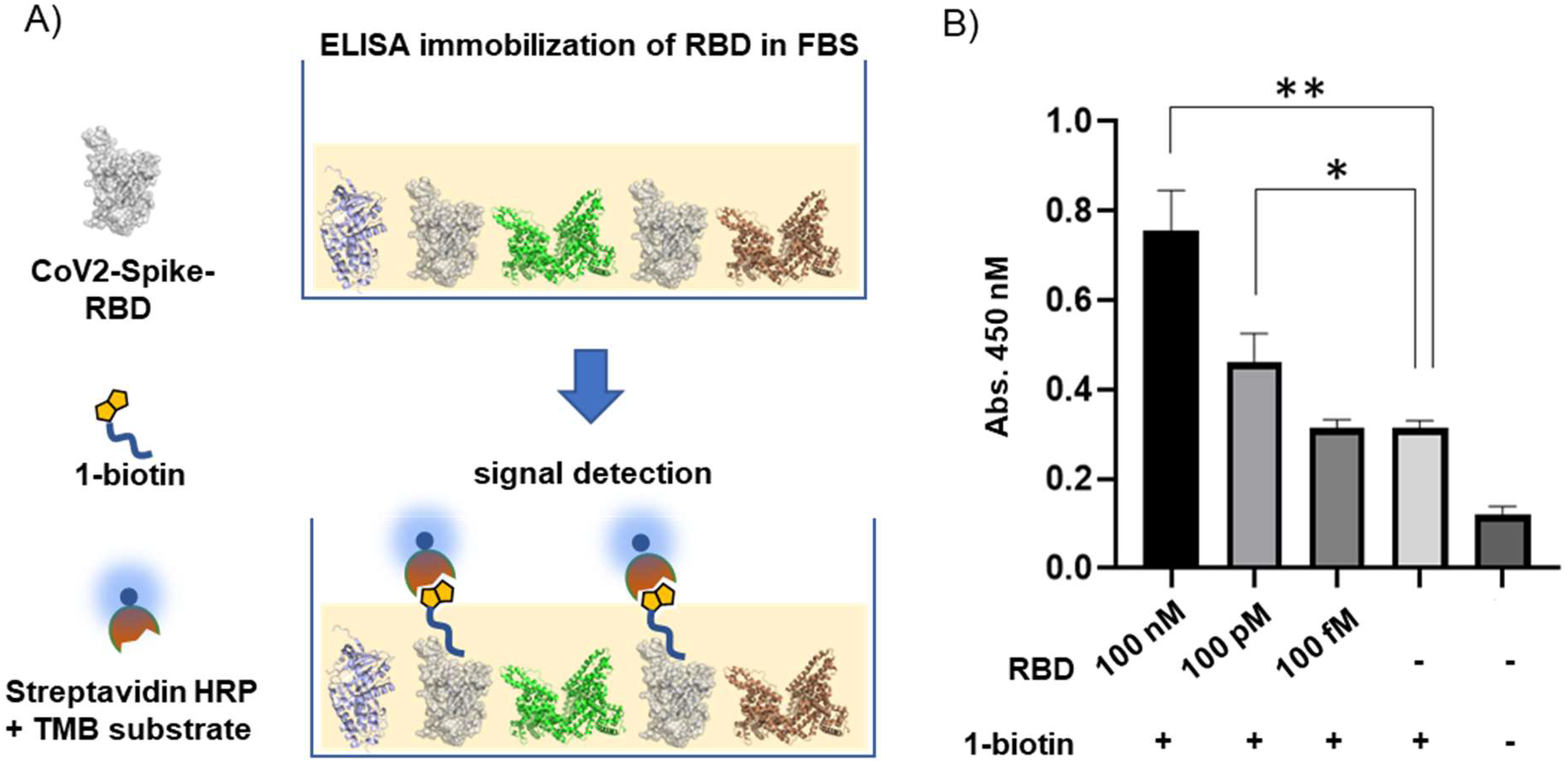
Picomolar SARS-CoV-2-spike-RBD quantities were detected by ELISA. A) Schematic representation of the ELISA assay: RBD dissolved in fetal bovine serum (FBS) in serial dilutions (100 nM – 100 fM) was immobilized on an ELISA plate. The plate was incubated with **1-biotin**, followed by streptavidin-HRP and TMB substrate. B) ELISA absorbance readout at 450 nM. Measurements were performed in technical triplicates (*n = 3*), and statistical significance was calculated with the unpaired t-test. RBD 100 nM vs no RBD: *p = 0.0012 (**)*; RBD 100 pM vs no RBD: *p = 0.018 (*)*.

## Concluding Remarks

Taken together, we have shown the use of our affinity selection – mass spectrometry platform^27^ for the rapid discovery of peptides binding to the SARS-CoV-2-spike-RBD. From 800 million synthetic peptides screened, we identified three sequences with a shared residue motif and nanomolar affinity. These peptides bind SARS-CoV-2-spike-RBD with selectivity over multiple human serum proteins and could detect it at picomolar concentrations in an ELISA format. Cross-binding of peptide **1** to the MERS coronavirus spike protein indicated a possible binding site distal from the binding site for the human ACE2 receptor.

The peptides reported here are potential starting points for the development of affinity-based diagnostic tools.^32–34^ High-affinity reagents without direct competition activity for native receptors could be used for virus-directed delivery of antiviral payloads, or for the development of proteasome or lysosome targeting chimeras (PROTACs^35^ and LYTACs^36^). Adapting peptide **1** to a chemiluminescence enzyme immunoassay or a similar assay could improve its sensitivity for detection of SARS-CoV-2.^8,37^ Because of its selectivity in biological media, peptide **1** could be utilized for the direct detection of SARS-CoV-2 in bodily fluids. Improved diagnostics are a topic of intense COVID-19 research as they may provide rapid, reliable and early detection.^38^ While direct detection suffers from low sensitivity,^10,39^ the rapid identification of SARS-CoV-2 is critical for patient contact tracing, identifying hosts, and epidemiologic studies.^2–4^ The peptides discovered by our platform may provide a useful SARS-CoV-2 detection modality to help achieve these goals.

## Notes

B.L.P. is a co-founder of Amide Technologies and Resolute Bio. Both companies focus on the development of protein and peptide therapeutics.

## ACKNOWLEDGMENTS

This research was supported by a COVID-19 Fast Grant award sponsored by Emergent Ventures at the Mercatus Center, George Mason University, and by a Bristol-Myers Squibb Unrestricted Grant in Synthetic Organic Chemistry awarded to B.L.P. S.P. thanks the Deutsche Forschungsgemeinschaft for a postdoctoral fellowship (DFG, PO 2413/1-1). Y.-C.L. is supported by the Deutsche Forschungsgemeinschaft (DFG, LE 4224/1-1). M.J. gratefully acknowledges postdoctoral fellowship support from the Rothschild Foundation, the Fulbright Program, and the Israel Council for Higher Education (VATAT). We thank the Biophysical Instrumentation Facility at MIT for providing access to the Octet Red96 Bio-Layer Interferometry System (NIH S10OD016326). We gratefully acknowledge Andrei Loas for helping with the preparation of the manuscript and organizational support during the project.

