## Supporting Information for "*De Novo* Discovery of High Affinity Peptide Binders for the SARS-CoV-2 Spike Protein"

### General information

**Chemicals**: Unless otherwise noted, all chemicals were obtained from commercial sources and used as received without further purification. Tetrahydrofuran (THF) and *N*,*N*-diisopropylethylamine (DIEA) were obtained from a Seca Solvent Purification System by Pure Process Technology.

**Materials for peptide synthesis:** H-Rink Amide-ChemMatrix resin was purchased from PCAS BioMatrix Inc. (St-Jean-sur-Richelieu, Quebec, Canada). 30 µm TentaGel M NH2 microspheres (M30352; 0.20 to 0.25 mmol/g amine loading) were purchased from Rapp Polymere (Tübingen, Germany). 20 µm TentaGel S NH2 microspheres (TMN-9909-PI; 0.2 to 0.3 mmol/g amine loading) was purchased from Peptides International (Louisville, KY). Fmoc-Ala-OH, Fmoc-Arg(Pbf)-OH, Fmoc-Asn(Trt)-OH, Fmoc-Asp(tBu)-OH, Fmoc-Gln(Trt)-OH, Fmoc-Glu(tBu)-OH, Fmoc-Gly-OH, Fmoc-His(Trt)-OH, Fmoc-Leu-OH, Fmoc-Lys(Boc)-OH, Fmoc-Met-OH, Fmoc-Phe-OH, Fmoc-Pro-OH, Fmoc-Ser(tBu)-OH, Fmoc-Thr(tBu)-OH, Fmoc-Trp(Boc)-OH, Fmoc-Tyr(tBu)-OH, and Fmoc-Val-OH were purchased from Advanced ChemTech (Louisville, KY). Trifluoroacetic acid (TFA; for HPLC, ≥99%), piperidine (ReagentPlus; 99%), triisopropylsilane (98%), 1,2-ethanedithiol (≥98%), were purchased from MilliporeSigma (St. Louis, MO). Diisopropylethylamine (99.5%; biotech. grade; DIEA) was also purchased from MilliporeSigma, and purified by passage through an activated alumina column (Pure Process Technology solvent purification system; Nashua, NH). Water was deionized using a Milli-Q Reference water purification system (Millipore).

**Materials for affinity selection:** Mouse anti-hemagglutinin (HA) monoclonal antibody clone 12ca5 (anti-HA mAb 12ca5) was purchased from Columbia Biosciences (Frederick, MD). SARS-CoV-2 RBD was purchased from Sino Biological (40592-V08H) and biotinylated in house (see: procedure for SARS-CoV-2 RBD biotinylation). HyClone™ Fetal Bovine Serum (SH30071.03HI, heat inactivated) was purchased from GE Healthcare Life Sciences (Logan, UT). Bovine serum albumin (BSA; RIA grade) and Tween 20 (reagent grade) were purchased from Amresco (Solon, OH). Dynabeads MyOne Streptavidin T1 magnetic microparticles were purchased from Invitrogen (Carlsbad, CA).

**LC-MS analysis**: LC-MS chromatograms and associated mass spectra were acquired using an Agilent 6550 ESI-QToF mass spectrometer. Mobile phases used for LC-MS analysis are solvent A (0.1% formic acid in water) and solvent B (0.1% formic acid in acetonitrile). The following LC-MS methods were used:

*Method A*: On 6550 MS; $C_4$ Phenomenex Jupiter column (1 x 150 mm, 5 µm); LC conditions: 1% B from 0–2 minutes, linear ramp from 5% to 61% B from 2–12 minutes, 0.1 mL/min flow rate.

*Method B*: On 6520 MS; Zorbax 300SB-$C_3$ column (2.1 x 150 mm, 5 µm); LC conditions: 1% B from 0–2 min, linear ramp from 5% to 61% B from 2–12 min, linear ramp from 61% to 90% 11-12 min, 0.8 mL/min flow rate.

**Purification**: Reversed-phase column chromatography was performed with a Biotage Selekt flash purification system equipped with 25 g Biotage Sfär Bio C18 D 20 µm columns with an appropriate gradient of MeCN/$H_2O$ (0.1% TFA).

Flash gradient: 15% MeCN, 1 column volume (CV); 15-60% MeCN, 8 CV; 60-95% MeCN 1 CV, 95% MeCN, 3 CV.

### Peptide synthesis procedures

**General procedure for manual solid-phase peptide synthesis (SPPS)**. ChemMatrix® Rink amide resin (loading 0.49 mmol/g, typical scale: 200 mg, 0.1 mmol) was loaded into a fritted syringe (12 mL), swollen in DMF (5 mL) for 5 minutes and then drained. Each Nα-Fmoc protected amino acid (1.0 mmol, 10 equiv) was dissolved in DMF containing 0.39 M HATU (2.5 mL). Immediately before the coupling DIEA (500 µL, 30 equiv) was added to the mixture to activate the amino acid. This solution after 15 seconds was added to the resin and reacted for 10 min, with occasional stirring. After completion of the coupling step, the syringe was drained, and the resin was washed with DMF (3 x 10 mL). Fmoc deprotection was performed by addition of piperidine (20% in DMF, 5 mL), to the resin (1 x 1 min + 1 x 5 min), followed by draining and washing the resin with DMF (5 x 10 mL). Side chain protection was as follows: Asn(Trt), Asp(O$^t$Bu), Cys(Trt), Gln(Trt), Glu(O$^t$Bu), His(Trt), Lys(Boc), Ser($^t$Bu), Thr($^t$Bu), Trp(Boc), Tyr($^t$Bu).

**Procedure for AFPS**: Peptides 1-biotin, 2-biotin, 4-biotin, scrambled1-1-biotin and scrambled2-1-biotin were synthesized on an automated flow synthesizer according to Hartrampf et al.[1] and Mijalis et al.[2]

**Procedure for peptide biotinylation:** All Lys-biotin residues were manually coupled to the C-terminus (see manual SPPS). For peptides synthesized by AFPS, Fmoc deprotection of the C-terminal Lys-biotin was performed in flow.

**Procedure for peptide cleavage**. Upon completion of the peptide synthesis, the resin was treated with a cleavage cocktail containing 94% TFA, 2.5% water, 2.5% 1,2-ethanedithiol (EDT) and 1% triisopropylsilane (TIPS) (v/v) at room temperature for 2 h. The TFA volume was then reduced under $N_2$ stream and cold diethyl ether (–80 °C) was added to precipitate the peptide. The resulting suspension was centrifugated at 4000 rpm for 3 min and the liquid was discarded. After repeating this step twice more, the pellet was dissolved in Milli-Q water with 0.1% TFA and lyophilized.

### Procedure for SARS-CoV-2-Spike-RBD biotinylation

EZ-Link-Sulfo-NHS-LC-LC-biotin (10 mM in $H_2O$, 15 µL, 150 nmol) was added to SARS-CoV-2-Spike-RBD (16 µM in PBS, pH = 7.5, 2 mL, 33 nmol) at 0 °C and then shaken on a nutating mixer for 2 h at ambient temperature. Reaction was quenched with addition of 1M Tris (50 µL). The excess biotin was removed from the mixture by size exclusion centrifugation through 10 kDa Amicon tubes (3 x 5 mL PBS). Concentration of biotinylated SARS-CoV-2-Spike-RBD was measured by absorption at 280 nm.

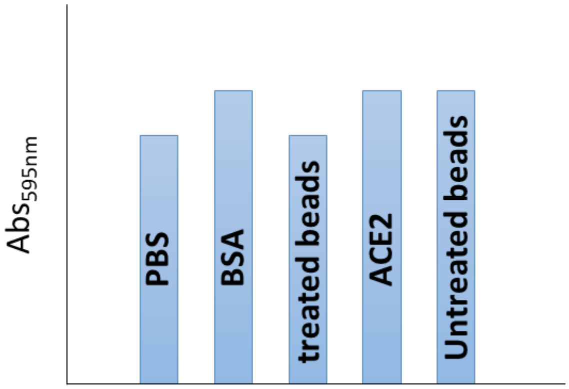

**SI Figure 1. Biotinylated SARS-CoV-2-RBD was immobilized on beads and structural integrity tested by ACE2 capture.** Successful biotinylation and integrity of the RBD and in particular ACE2 binding site after reaction were confirmed as followed: MyOne streptavidin beads (100 µL, 10 mg/mL) were washed, suspended in PBS buffer (1 mL, pH = 7.5) and Biotin-SARS-CoV-2-Spike-RBD was added (50 µL, 18 µM). The mixture was incubated at 4 °C for 20 minutes to immobilize RBD on the beads. Then the beads were washed with PBS (3 x 1 mL) and incubated with ACE2 (100 µL, 0.025mg/mL) for 30 minutes at 4 °C. Also, beads without SARS-CoV-2-Spike-RBD were treated with the same amount of ACE2, as control. The supernatant of the two mixtures were collected and protein quantity was determined via Bradford assay (Bio Rad, according to vendor instructions). Additional controls were: PBS, ACE2 (0.025 mg/mL) and bovine serum albumin (0.025 mg/mL). Final colorimetric readout (at 595 nM) confirmed the quantitative depletion of ACE2 from mixture incubated with SARS-CoV-2-Spike-RBD-beads.

Biotin-SARS-CoV-2-Spike-RBD was stored at -80 °C.

### Affinity selection and mass spectrometry

According to Quartararo et al, *Nat. Comm.*, **2020**[3]

**Preparation of SARS-CoV-2-Spike-RBD and 12ca5-functionalized magnetic beads:** MyOne Streptavidin T1 Dynabeads (300 µL of 10 mg/mL stock) were transferred to 1.7 mL plastic centrifuge tubes, and placed in a magnetic separation rack. Beads were washed 3 x 1 mL w/ 10% FBS, 0.02% Tween 20, 1x PBS, and then treated with 44 µL of biotinylated SARS-CoV-2-Spike-RBD (18 µM) or 347 µL 12ca5 (1.5 µM; 0.45 nmol). The resulting suspensions were transferred to a rotating vertical mixer and allowed to incubate for 30 min at 4°C. After this time, the beads were returned to the separating rack, the supernatant was removed, and the beads were washed 3 x 1 mL w/ 10% FBS, 0.02% Tween 20, 1x PBS. Beads were resuspended in 300 µL of 10% FBS, 0.02% Tween 20, 1x PBS.

**Affinity capture:** Library (10 fmol/member) was incubated with 100 µL (1 mg) portions of protein-immobilized magnetic beads (prepared above) in the presence of 10% FBS, 1x PBS (final volume: 1 mL) on a rotating mixer for 1 h at 4 °C. Final conditions: 1 mg/mL magnetic beads, 10 pM/member library.

**Elution:** The centrifuge tubes containing the bead suspensions were transferred to the magnetic separation rack. The beads were washed 3 x 1 mL w/ 1x PBS. Bound peptides were eluted with 2 x 100 µL 6M guanidine hydrochloride, 200 mM phosphate, pH 6.8. Eluates were concentrated via C18 ZipTip® pipette tips (according to vendor protocol) and lyophilized.

**NanoLC-MS/MS analysis:** The standard nano-LC method was run at 40 °C and a flow rate of 300 nL/min with the following gradient: 1% solvent B in solvent A ramping linearly to 41% B in A over 120 min, where solvent A = water (0.1% FA), and solvent B = 80% acetonitrile, 20% water (0.1% FA). Positive ion spray voltage was set to 2200 V. Orbitrap detection was used for primary MS with the following parameters: resolution = 120,000; quadrupole isolation; scan range = 200-1400 m/z; RF lens = 30%; AGC target = $1 \times 10^6$; maximum injection time = 100 ms; 1 microscan.

Acquisition of secondary MS spectra was done in a data-dependent manner: dynamic exclusion was employed such that a precursor was excluded for 30 s if it was detected four or more times within 30 s (mass tolerance: 10.00 ppm); monoisotopic precursor selection used to select for peptides; intensity threshold was set to $5 \times 10^4$; charge states 2-10 were selected; and precursor selection range was set to 200-1400 m/z. The top 15 most intense precursors that met the preceding criteria were subjected to subsequent fragmentation.

Three fragmentation modes – collision-induced dissociation (CID), higher-energy collisional dissociation (HCD), and electron-transfer/higher-energy collisional dissociation (EThcD) – were used for acquisition of secondary MS spectra. Only precursors with charge states 3 and above were subjected to all three fragmentation modes; precursors with charge states of 2 were subjected to CID and HCD only. For all three modes, detection was performed in the Orbitrap (resolution = 30,000; quadrupole isolation; isolation window = 1.3 m/z; AGC target = $2 \times 10^4$; maximum injection time = 100 ms; 1 microscan). For CID, a collision energy of 30% was used. For HCD, a collision energy of 25% was used. For EthcD, a supplemental activation collision energy of 25% was used.

***De novo* peptide sequencing:** *De novo* peptide sequencing was performed by processing .raw files obtained from Orbitrap analysis using PEAKS Studio (version 8.5) from Bioinformatics Solutions Inc. (ON, Canada). HCD and CID scans were merged within a 0.2 minute and 0.02 Da window, mass precursor correction was used, and primary mass filtration was employed as appropriate. Auto *de novo* sequencing was performed using a 15 ppm precursor mass error and 0.02 Da fragment mass error, and with the following modifications: fixed C-term amidation (-0.98 Da) on lysine, and variable oxidation on methionine (+15.99 Da). 15 candidate sequences were obtained for each preprocessed scan. Post-*de novo* data analysis was performed as described in Vinogradov, A. V. *et. al*. Library design-facilitated high-throughput sequencing of synthetic peptide libraries. *ACS Comb. Sci.* **19** 694-701 (2017).

### Outcome affinity selections

Hit sequence **1:** TVFGLNVWKRYSK
ALC = 81%
m/z = 399.9814
z = +4
RT = 60.31
Mass = 1595.894

A)

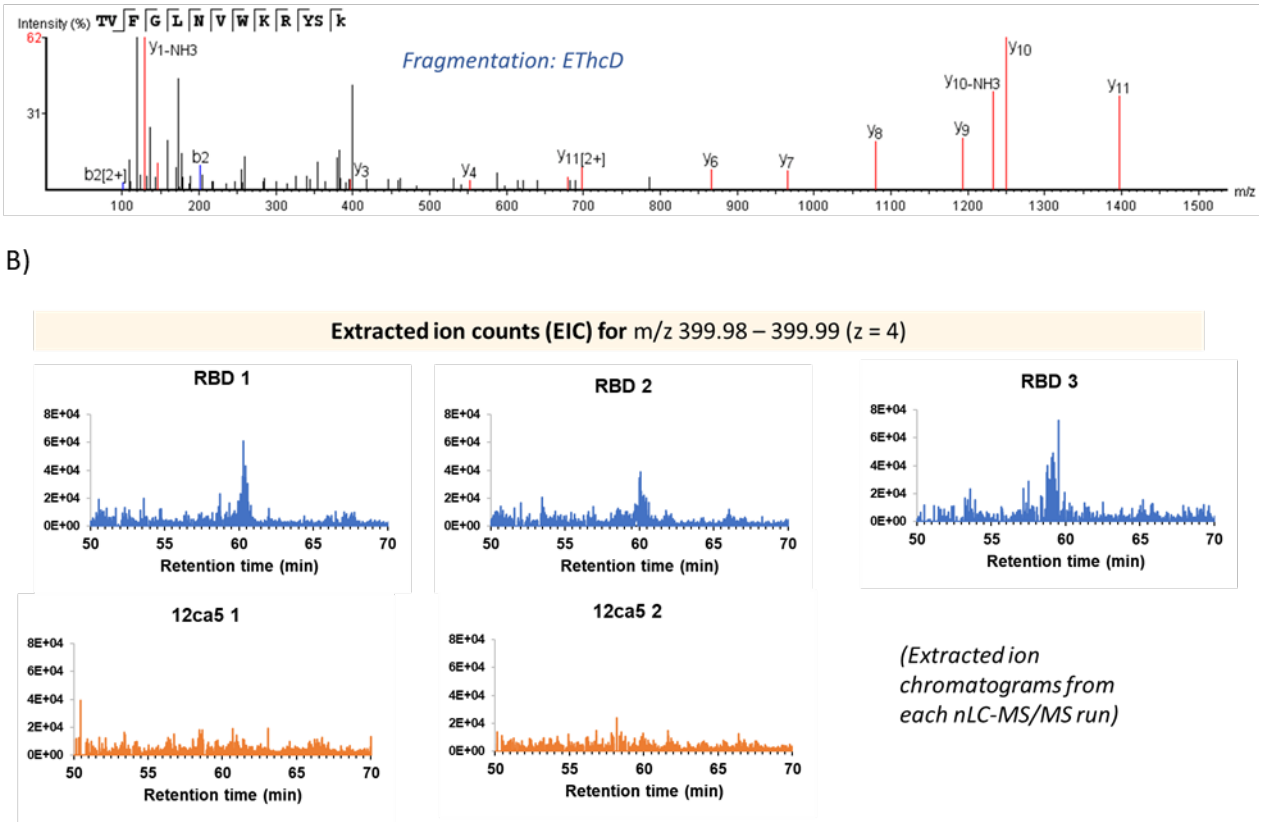

**SI Figure 2.** A) Electron transfer higher energy collision induced dissociation (EThcD) MS/MS spectrum of peptide 1 enriched by affinity selection. B) Extracted ion counts for precursor ion (m/z = 399.98-399.99) of enriched peptide 1 for SARS-CoV-2-spike-RBD samples (blue) and 12ca5 samples (orange).

Hit sequence **2:** LVOGLNAWNOWYK (O = methionine oxide)

ALC = 99%

m/z = 828.9074

z = +2

RT = 88.21

Mass = 1655.795

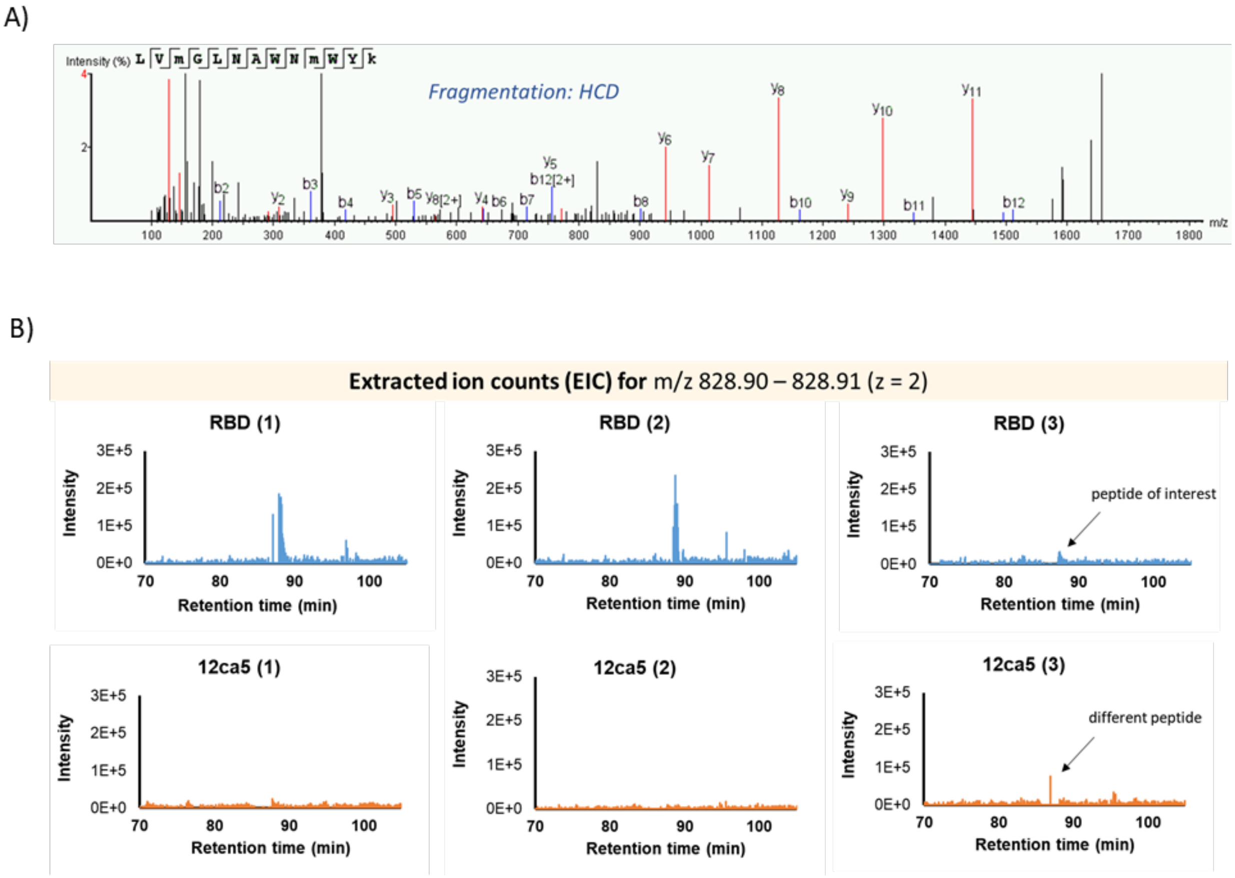

**SI Figure 3.** A) Higher energy collision induced dissociation (HCD) MS/MS spectrum of peptide 3 enriched by affinity selection. B) Extracted ion counts for precursor ion (m/z = 828.90-828.91) of enriched peptide 3 for SARS-CoV-2-spike-RBD samples (blue) and 12ca5 samples (orange).

Hit sequence **3:** LVOGLHVYLRQGK (O = methionine oxide)

ALC = 89%

m/z = 382.9759

z = +4

RT = 42.54

Mass = 1527.871

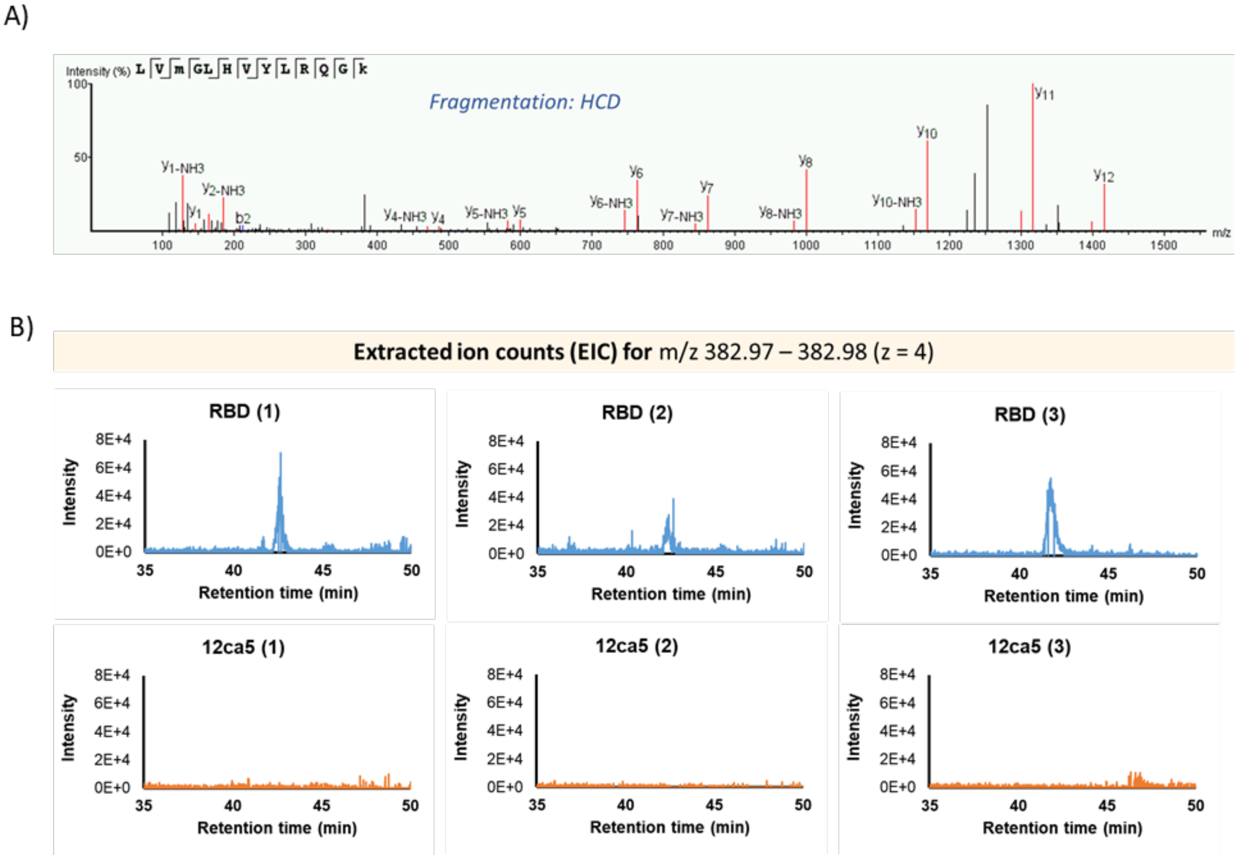

**SI Figure 4.** A) Higher energy collision induced dissociation (HCD) MS/MS spectrum of peptide 3 enriched by affinity selection. B) Extracted ion counts for precursor ion (m/z = 382.97-382.98) of enriched peptide 3 for SARS-CoV-2-spike-RBD samples (blue) and 12ca5 samples (orange).

LC-MS analysis of peptides

**1-Biotin: TVFGLNVWKRYSK-βA-K(Biotin)-CONH$_2$**

**AFPS synthesis, Biotage purification, flash gradient**

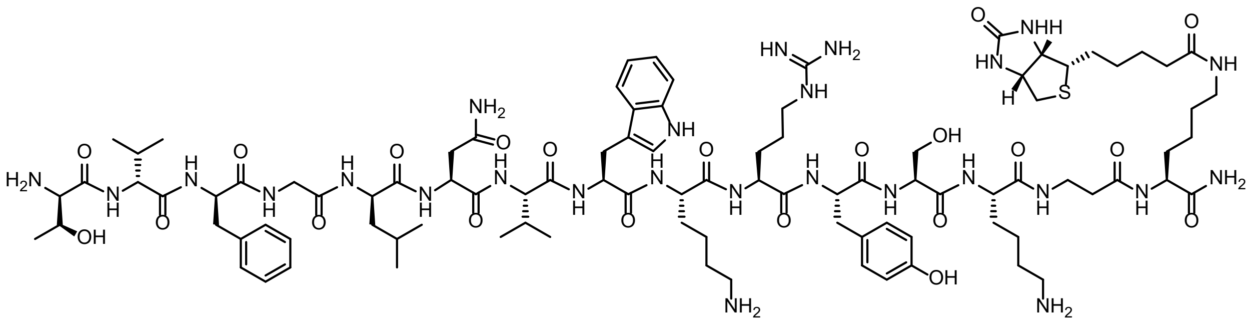

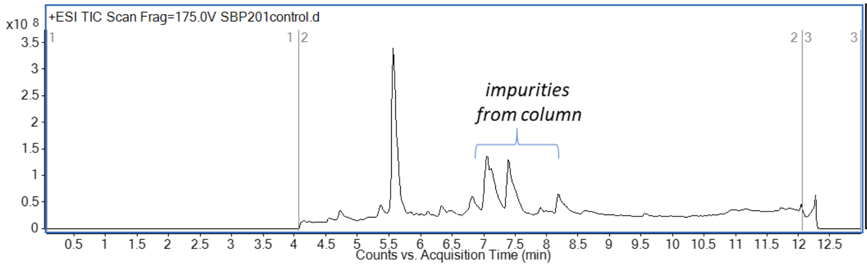

**HRMS** (SI-QToF) *m/z:* [M+H]$^+$
calcd. 2022.11
found 2022.11

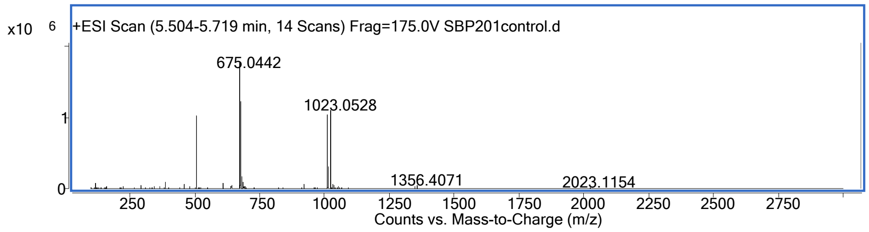

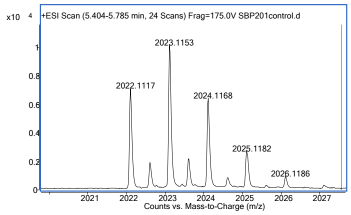

**2-Biotin: LVMGLNAWNMWYK-βA-K(Biotin)-CONH$_2$**

**AFPS synthesis, Biotage purification, flash gradient**

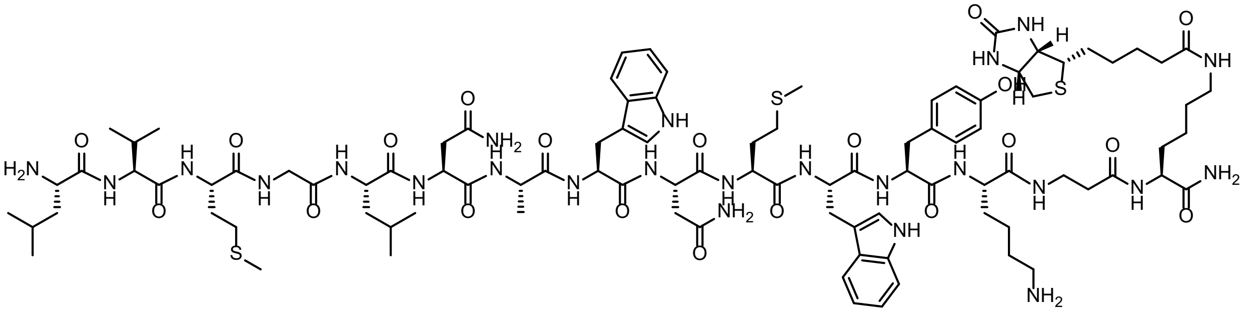

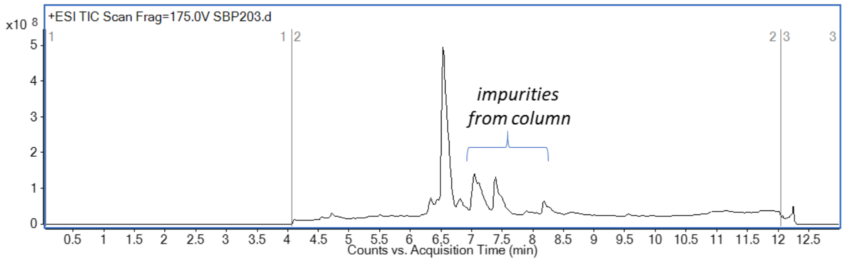

**HRMS** (SI-QToF) *m/z:* [M+H]$^+$
calcd. 2050.02
found 2050.03

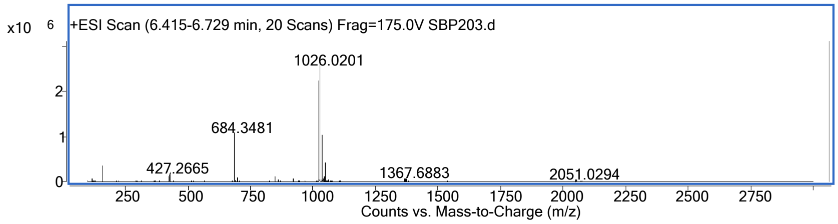

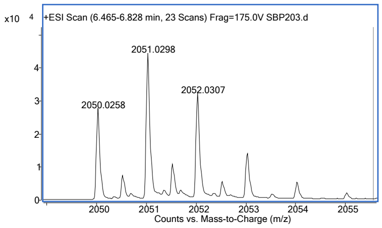

**3-Biotin: LVMGLHVYLRQGK-βA-K(Biotin)-CONH$_2$**

**AFPS synthesis, desalted via solid phase extraction**

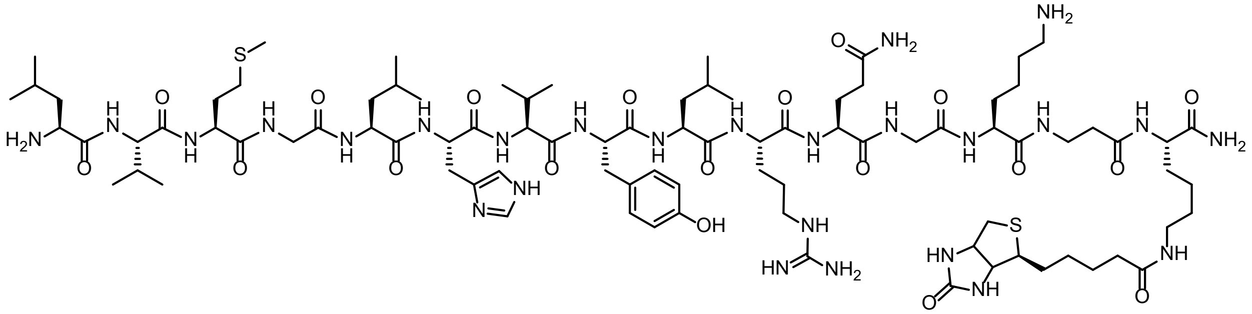

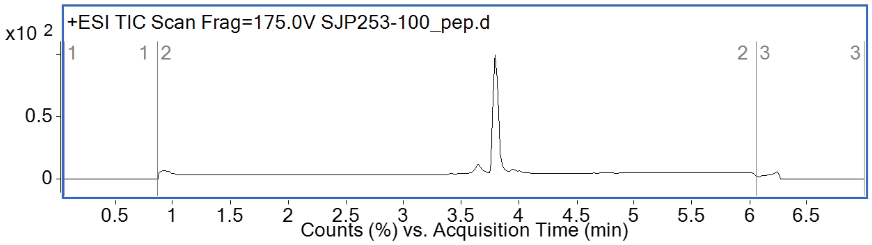

**HRMS** (SI-QToF) *m/z:* [M+H]$^+$
calcd. 1938.09
found 1938.09

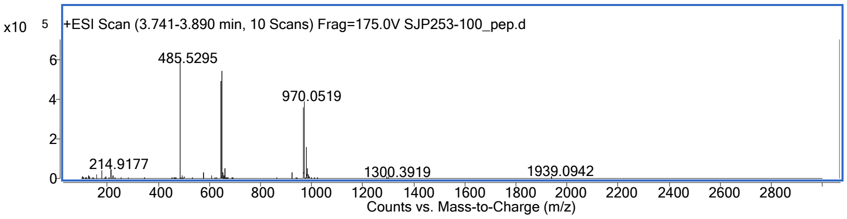

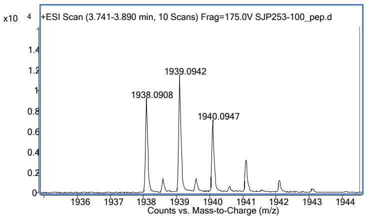

**4-Biotin: LVMGLNVWLRYSK-βA-K(Biotin)-CONH₂**

**AFPS synthesis, Biotage purification, flash gradient**

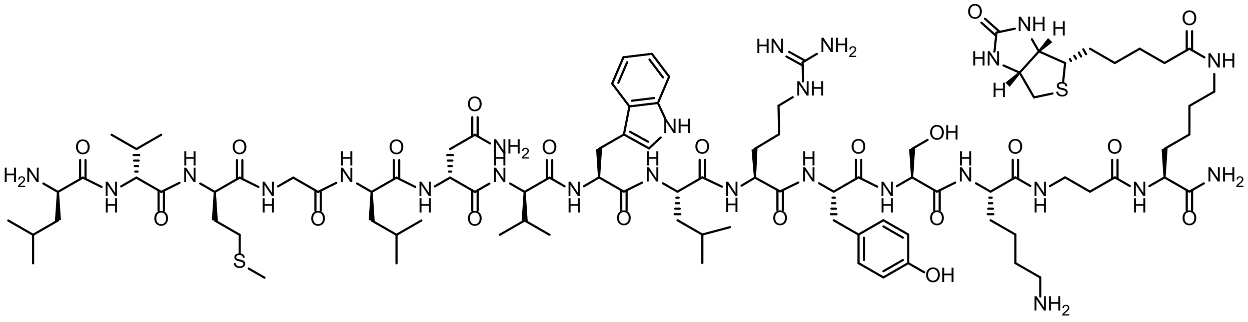

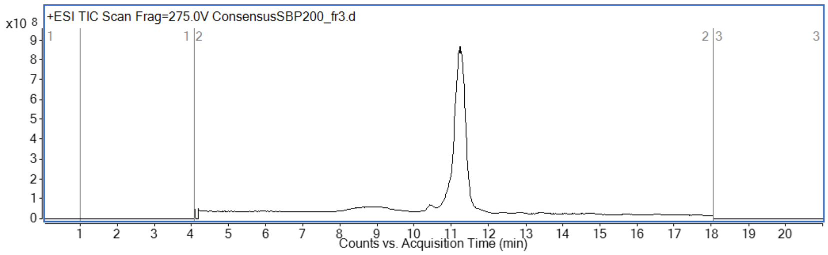

**HRMS** (SI-QToF) *m/z:* $[M+3H]^{3+}$
calcd. 668.373
found 668.373

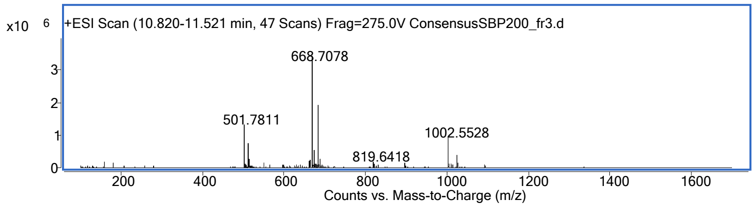

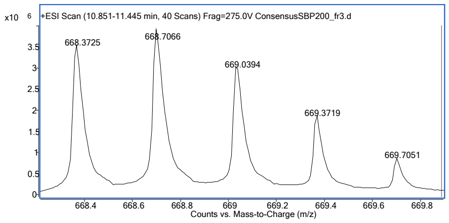

**Ac-1-Biotin: Ac-TVFGLNVWKRYSK-βA-K(Biotin)-CONH$_2$**

**Manual synthesis**

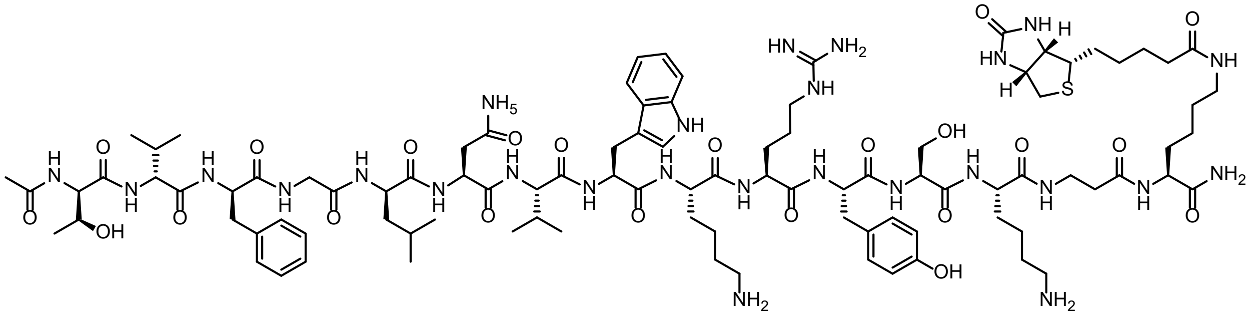

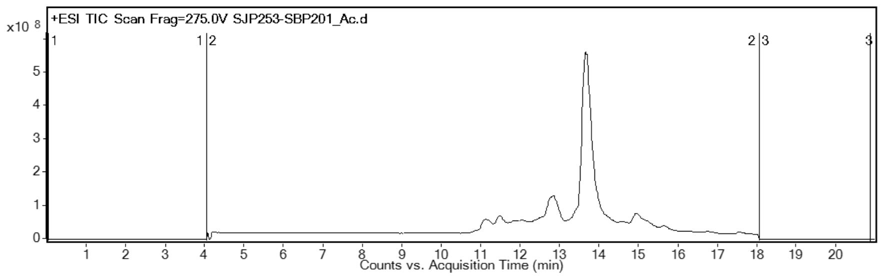

**HRMS** (SI-QToF) *m/z:* [M+3H]$^{3+}$
calcd. 688.713
found 688.718

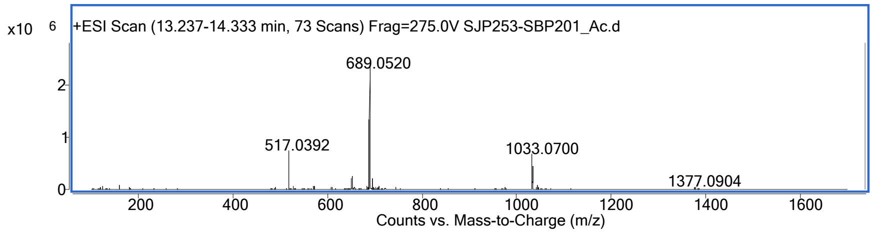

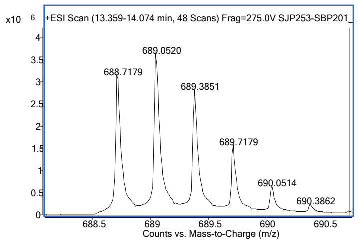

**1-Biotin-scrambled-1: GSVKRWLTYVKNF-βA-K(Biotin)-$CONH_2$**

**AFPS synthesis, Biotage purification, flash gradient**

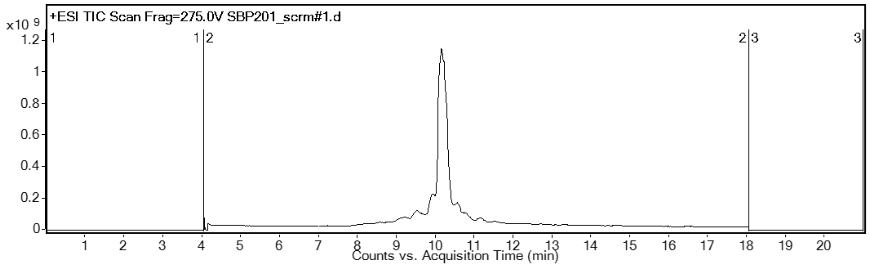

**HRMS** (SI-QToF) *m/z:* $[M+2H]^{2+}$
calcd. 1011.56
found 1011.56

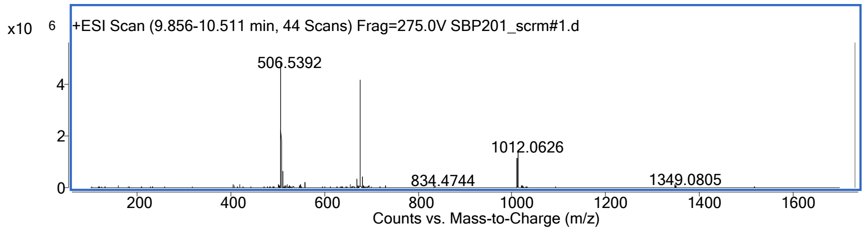

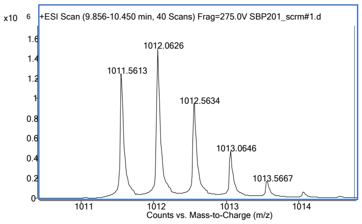

**1-Biotin-scrambled-2: RFYVTKGWSNKVL-βA-K(Biotin)-$CONH_2$**

**AFPS synthesis, Biotage purification, flash gradient**

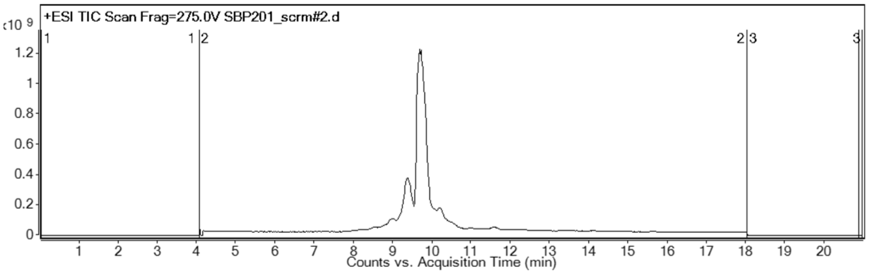

**HRMS** (SI-QToF) *m/z:* $[M+2H]^{2+}$
calcd. 1011.56
found 1011.56

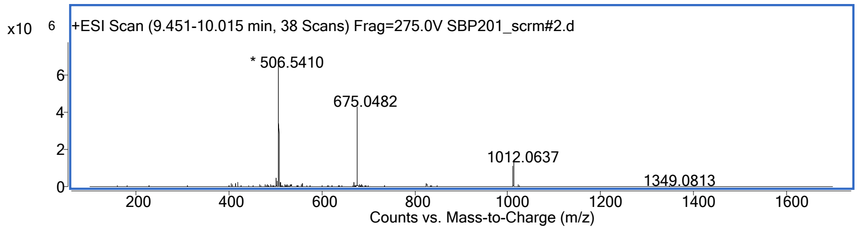

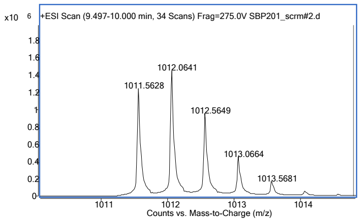

**Alanine scan of 1-Biotin: TVFGLNVWKRYSK-βA-K(Biotin)-$CONH_2$ (Manual SPPS, used without further purification after ether trituration and filtration)**

**AVFGLNVWKRYSK-βA-K(Biotin)-$CONH_2$**

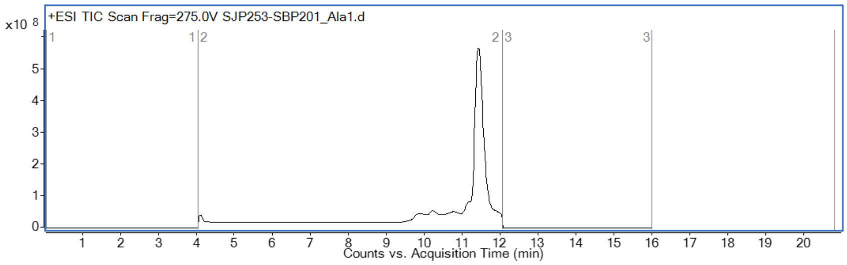

**HRMS** (SI-QToF) *m/z:* $[M+4H]^{4+}$
calcd. 498.780
found 498.785

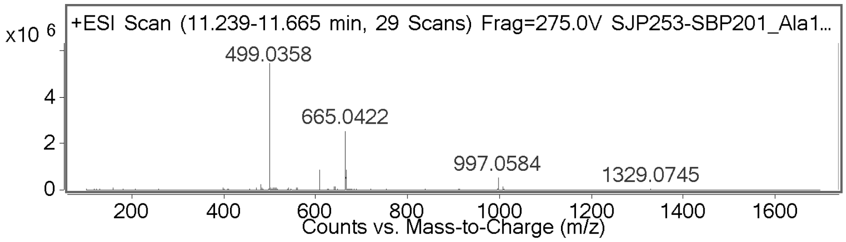

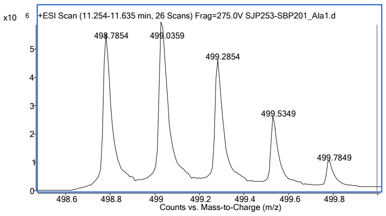

**TAFGLNVWKRYSK-βA-K(Biotin)-$CONH_2$**

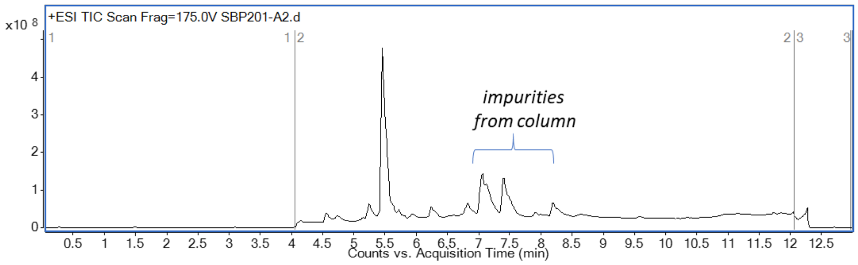

**HRMS** (SI-QToF) *m/z:* $[M+H]^{+}$
calcd. 1994.08
found 1994.08

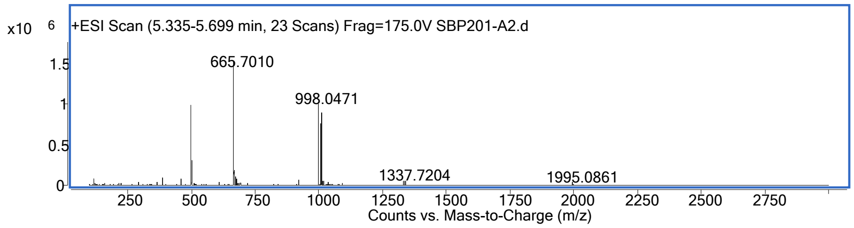

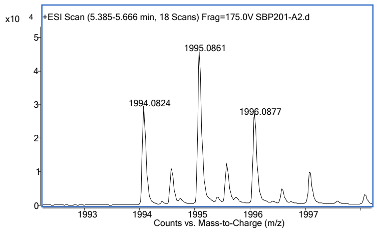

**TVAGLNVWKRYSK-βA-K(Biotin)-CONH$_2$**

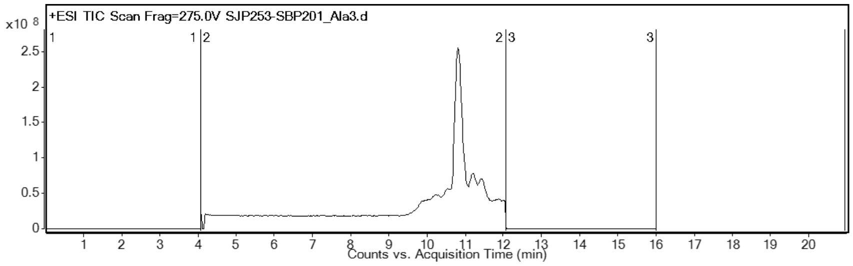

**HRMS** (SI-QToF) *m/z:* $[M+4H]^{4+}$
calcd. 487.275
found 487.279

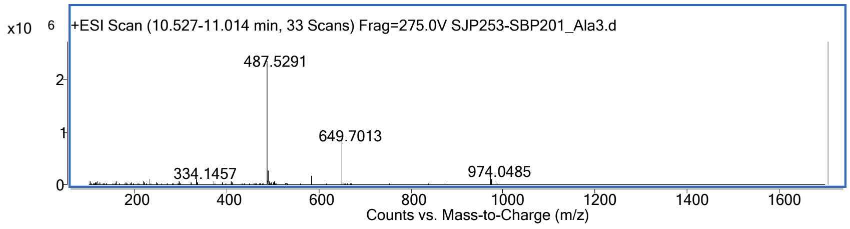

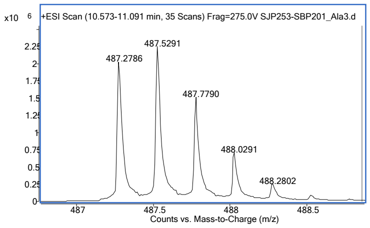

**TVFALNVWKRYSK-βA-K(Biotin)-CONH$_2$**

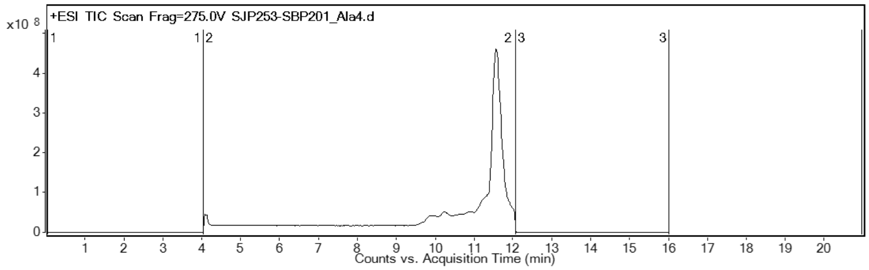

**HRMS** (SI-QToF) *m/z:* $[M+4H]^{4+}$
calcd. 509.787
found 509.792

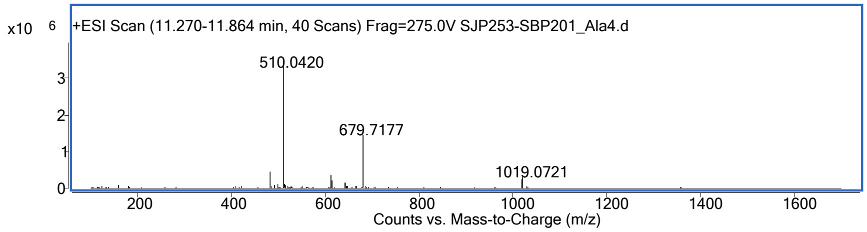

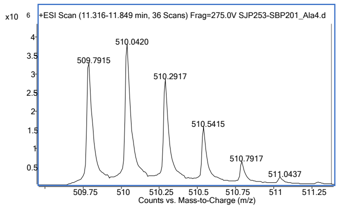

**TVFGANVWKRYSK-βA-K(Biotin)-CONH$_2$**

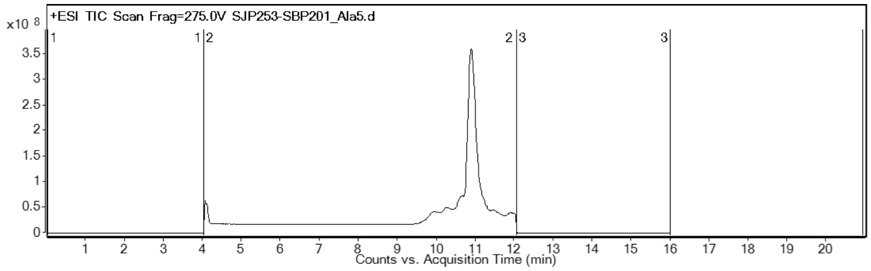

**HRMS** (SI-QToF) *m/z:* [M+4H]$^{4+}$
calcd. 496.772
found 496.775

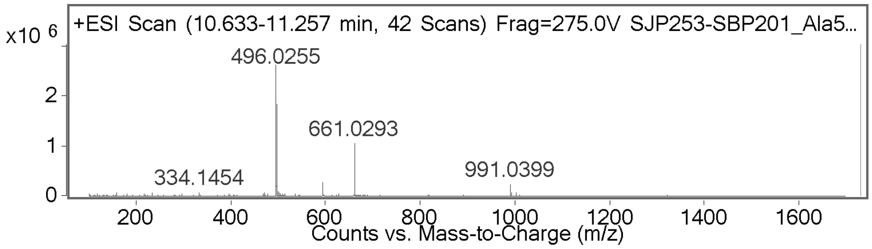

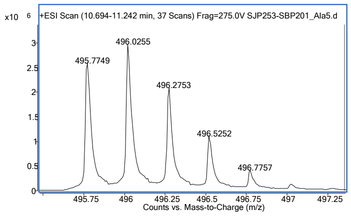

**TVFGLAVWKRYSK-βA-K(Biotin)-CONH$_2$**

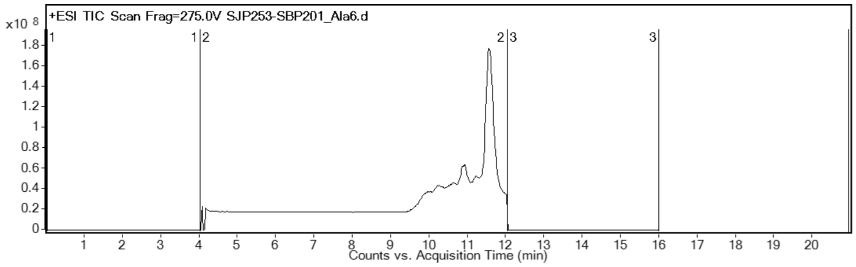

**HRMS** (SI-QToF) *m/z:* [M+4H]$^{4+}$
calcd. 495.533
found 495.534

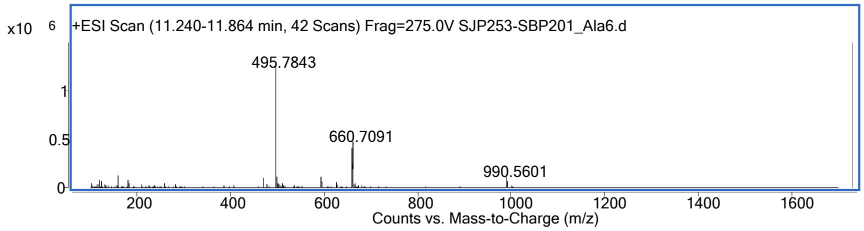

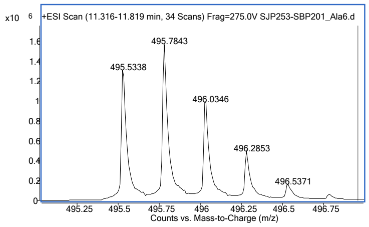

**TVFGLNAWKRYSK-βA-K(Biotin)-$CONH_2$**

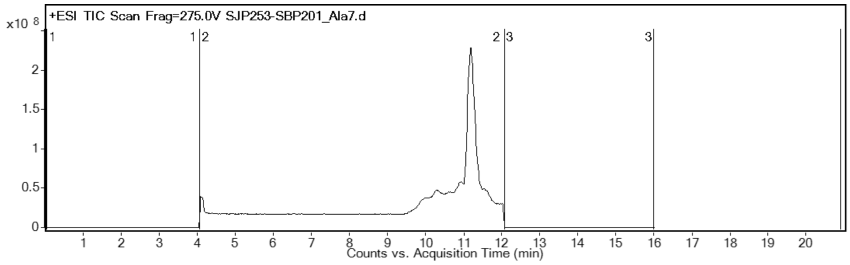

**HRMS** (SI-QToF) *m/z:* $[M+4H]^{4+}$
calcd. 499.275
found 499.278

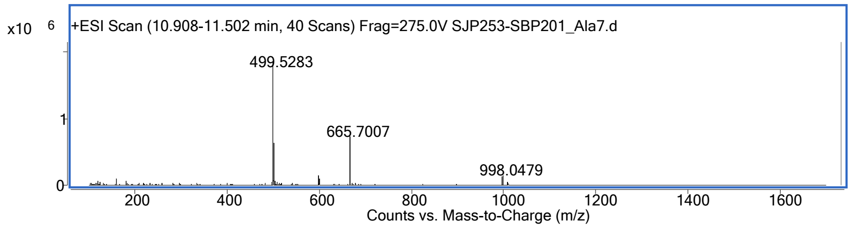

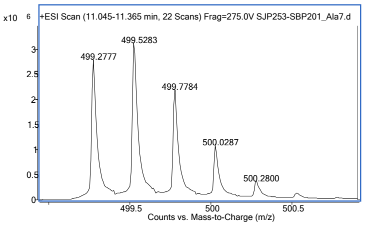

**TVFGLNVAKRYSK-βA-K(Biotin)-$CONH_2$**

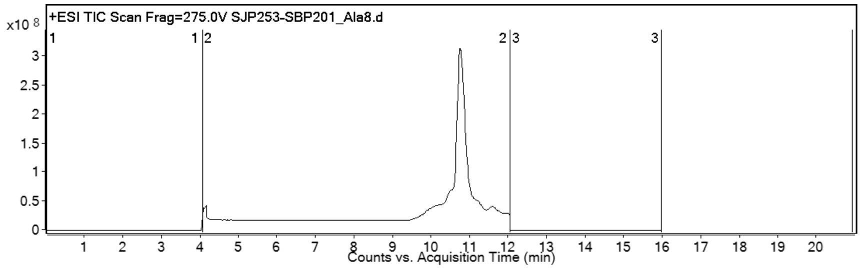

**HRMS** (SI-QToF) *m/z:* $[M+4H]^{4+}$
calcd. 477.523
found 477.525

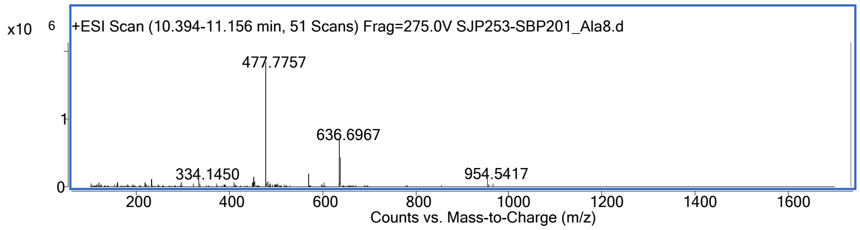

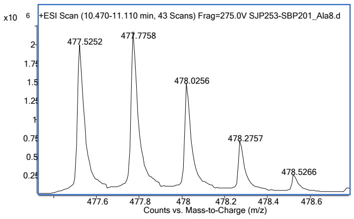

**TVFGLNVWARYSK-βA-K(Biotin)-CONH$_2$**

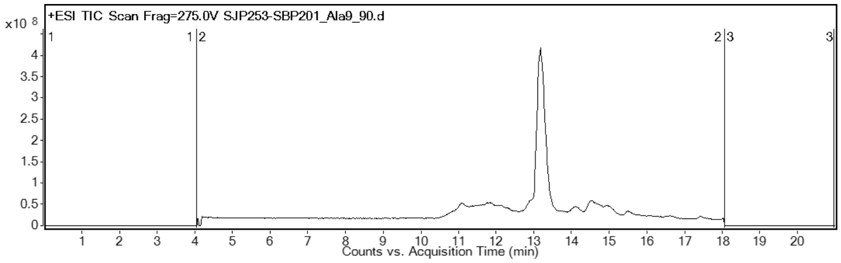

**HRMS** (SI-QToF) *m/z:* [M+3H]$^{3+}$
calcd. 655.691
found 655.693

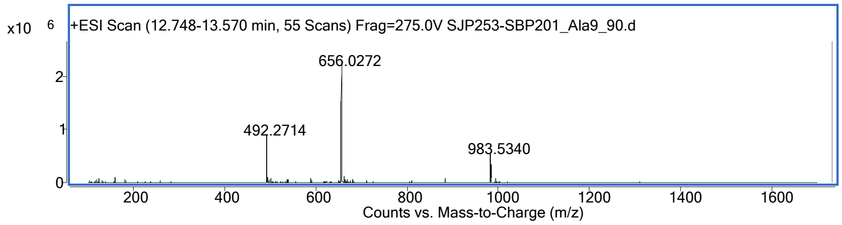

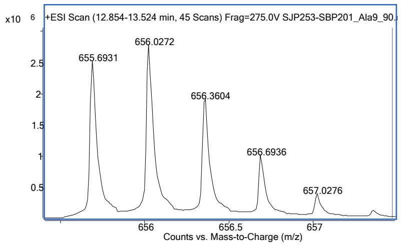

**TVFGLNVWKAYSK-βA-K(Biotin)-CONH$_2$**

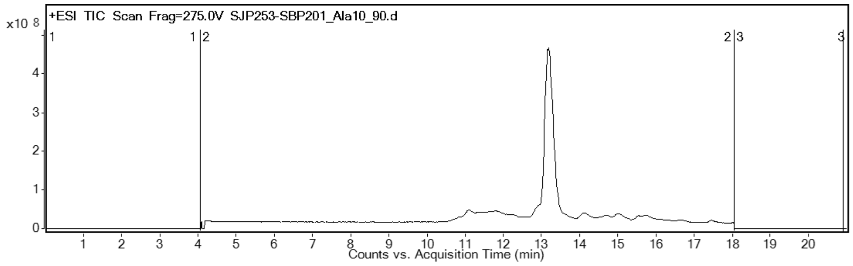

**HRMS** (SI-QToF) *m/z:* [M+3H]$^{3+}$
calcd. 646.354
found 646.357

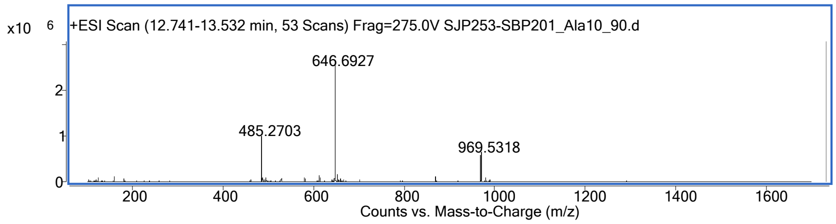

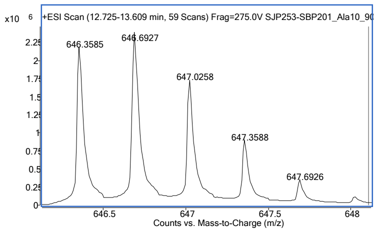

**TVFGLNVWKRASK-βA-K(Biotin)-$CONH_2$**

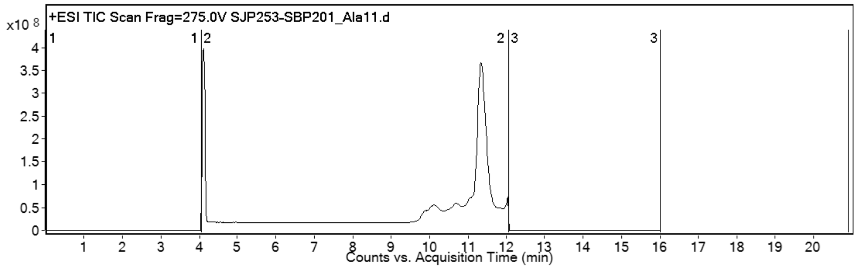

**HRMS** (SI-QToF) *m/z:* $[M+4H]^{4+}$
calcd. 483.275
found 483.277

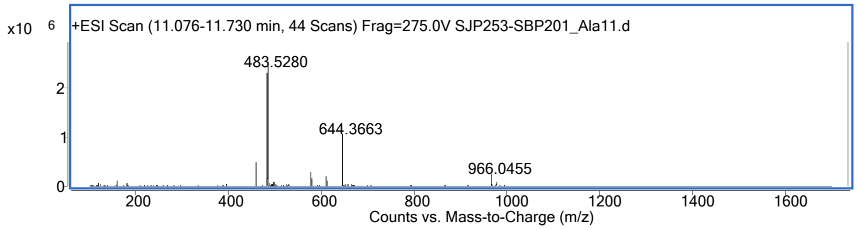

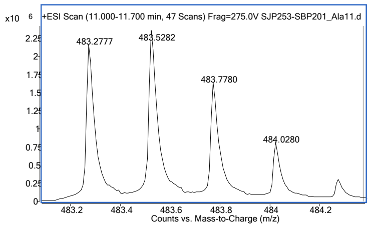

**TVFGLNVWKRYAK-βA-K(Biotin)-$CONH_2$**

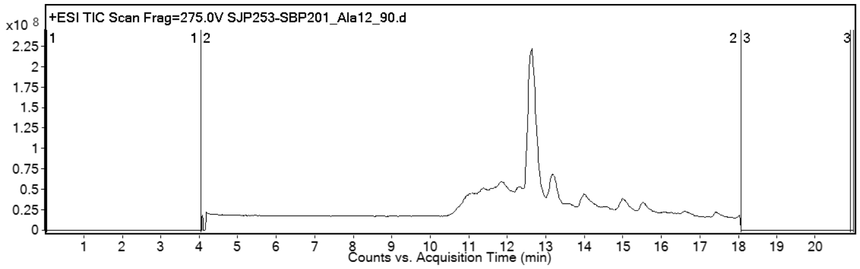

**HRMS** (SI-QToF) *m/z:* $[M+4H]^{4+}$
calcd. 502.285
found 502.287

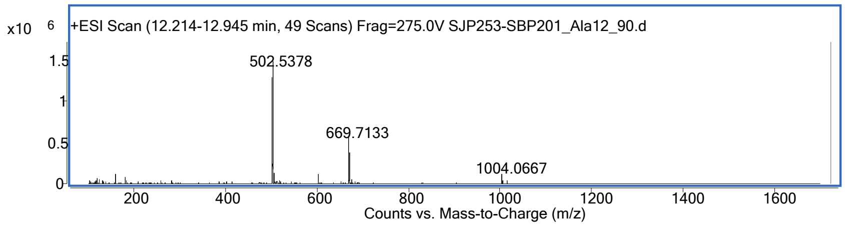

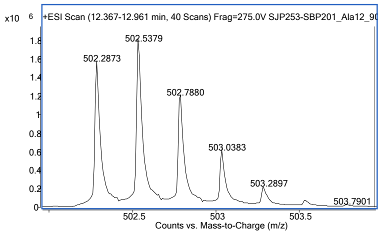

**Truncation scan of 1-Biotin: TVFGLNVWKRYSK-βA-K(Biotin)-CONH$_2$ (Manual SPPS, used without further purification after ether trituration and filtration)**

**-VFGLNVWKRYSK-βA-K(Biotin)-CONH$_2$**

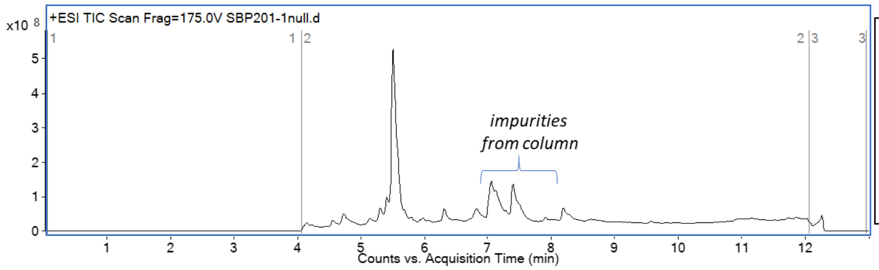

**HRMS** (SI-QToF) *m/z:* [M+H]$^+$
calcd. 1921.06
found 1921.07

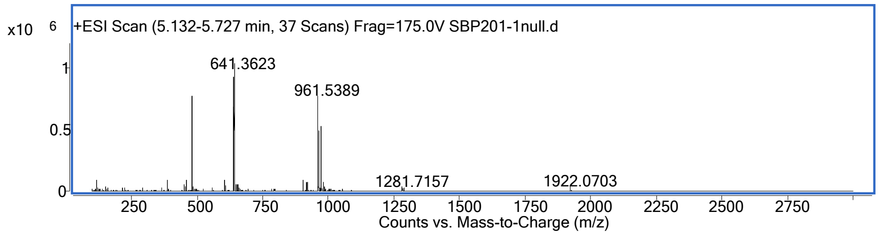

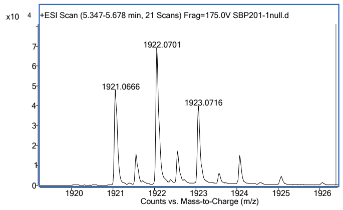

**--FGLNVWKRYSK-βA-K(Biotin)-CONH$_2$**

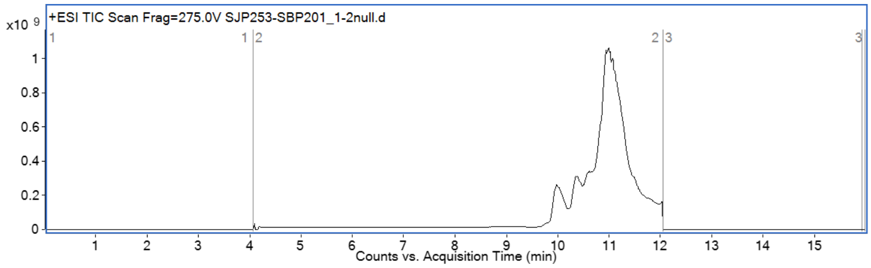

**HRMS** (SI-QToF) *m/z:* [M+4H]$^{4+}$
calcd. 456.255
found 456.262

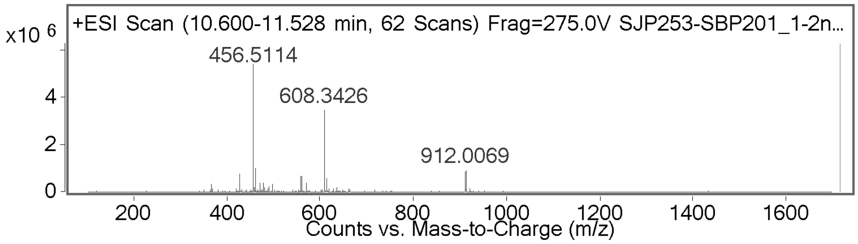

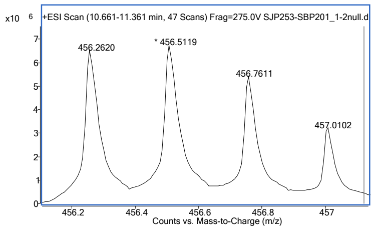

**TVFGLNVWKRY---βA-K(Biotin)-$CONH_2$**

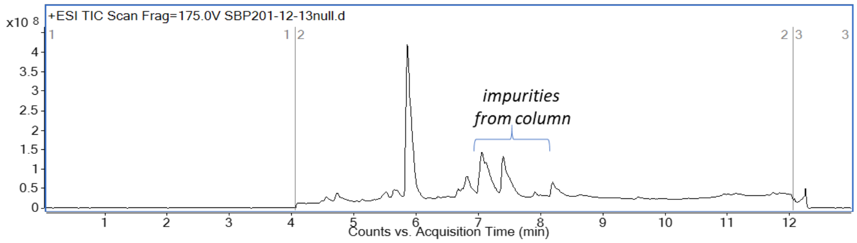

**HRMS** (SI-QToF) *m/z:* $[M+H]^+$
calcd. 1806.98
found 1806.99

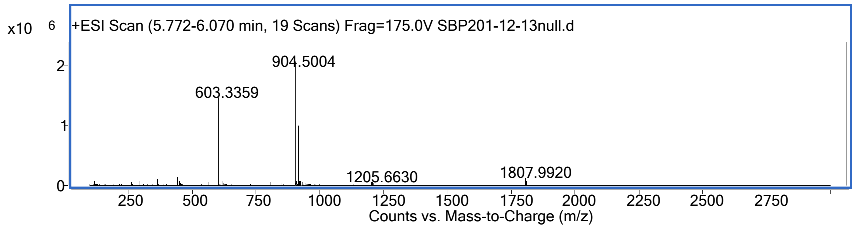

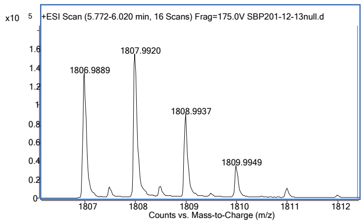

**TVFGLNVWKR----βA-K(Biotin)-$CONH_2$**

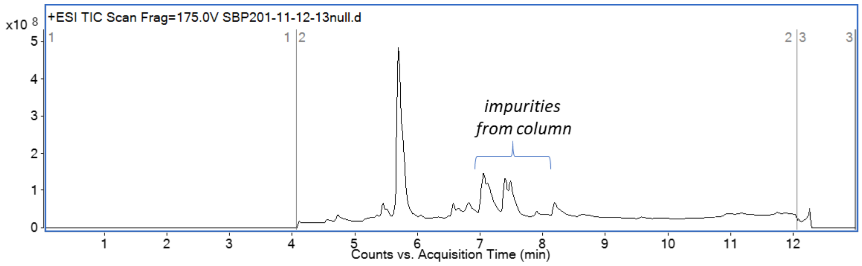

**HRMS** (SI-QToF) *m/z:* $[M+H]^+$
calcd. 1943.91
found 1643.93

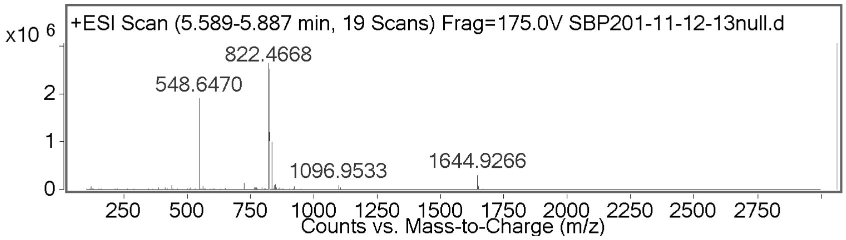

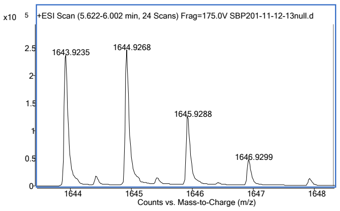

Biolayer interferometry (BLI)

#### General procedure

Lyophilized peptide biotinylated on C-term lysine) was dissolved to 2 mg/mL in 1x PBS and diluted 500-fold into 0.1% BSA, 0.02% Tween-20, 1x PBS ('kinetic buffer') for immobilization onto streptavidin Octet biosensors (ForteBio; Menlo Park, CA). Biolayer interferometry (BLI) assays were performed in 96 well plates (GreinerBio-One; Kremsmünster, Austria; polypropylene, flat-bottom, chimney well) using an Octet Red96 System (ForteBio; Menlo Park, CA). Wells were filled with 200 µL of kinetic buffer, peptide solution, or target protein solution (prepared in kinetic buffer at concentrations as indicated). Biotinylated peptide was immobilized onto the streptavidin tip for 120 s. Sensors were then dipped into kinetic buffer for 60 s, protein solutions 300 s, and finally into kinetic buffer for 300 s. Measurements were carried out at 30 °C. Data were analyzed within the ForteBio Data Analysis software. Data were processed by subtracting reference ligand only reference wells and aligning the data to the beginning of the association. Then, global kinetic fit was performed of all sensogram curves with a 1:1 model (Rmax linked). Steady state Kds are reported.

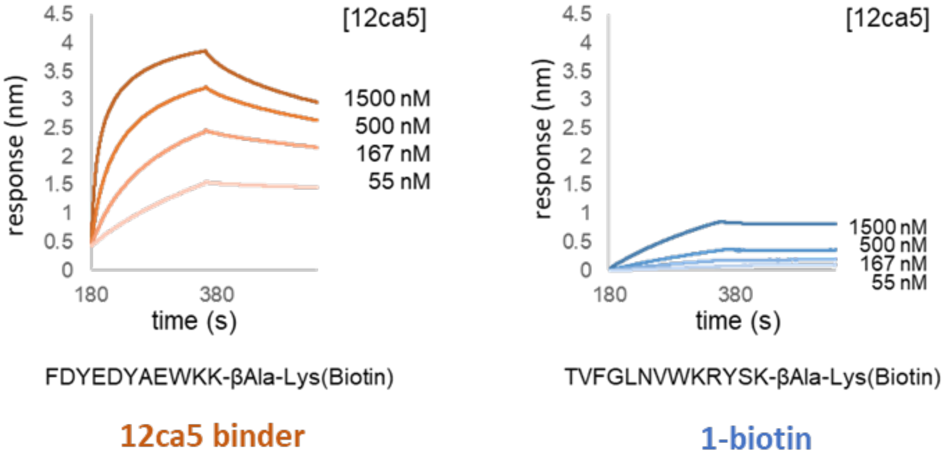

**SI Figure 5. Peptide 1 showed minimal association to 12ca5 compared to a known positive control 12ca5 binder.** Side to side analysis of binding to 12ca5 of a peptide selected for 12ca5 binding (orange line) and hit peptide **1** (blue line). Both peptides were immobilized on BLI tips and dipped into serial dilutions of 12ca5 for determination of association/dissociation

**Competition BLI**

**1-Biotin** was immobilized onto streptavidin Octet biosensors (according to general BLI procedure) and dipped into solution containing SARS-CoV2 RBD (500 nM) and peptide **1** (0, 1 µM, 2 µM, 4 µM, 8 µM and 16 µM). The maximum association response after 300 seconds was determined.

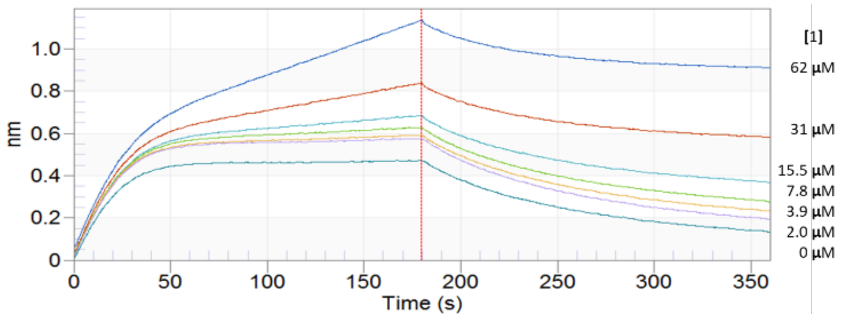

**SI Figure 6. Peptide hit 1 does not compete for the ACE2 binding site on the SARS-CoV-2-spike RBD.** Biotin-ACE2 was immobilized on BLI tips and dipped into solution containing a fixed concentration of SARS-CoV-2-spike RBD (100 nM) and different concentrations of peptide **1**. Assay conditions according Zhang et al.[4]

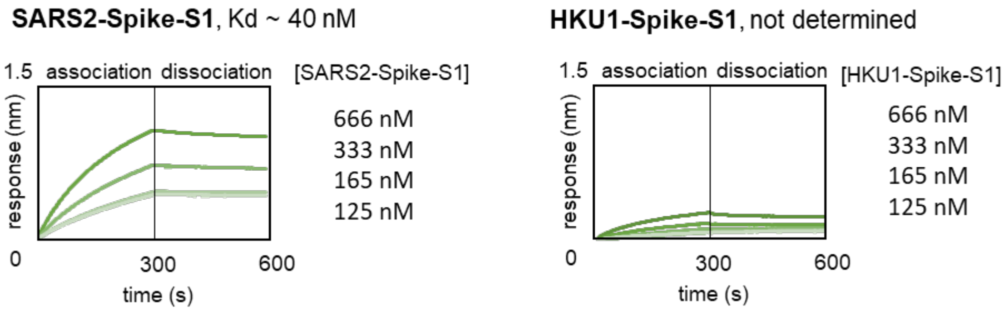

**SI Figure 7. BLI analysis of peptide 1 association to SARS-CoV-2-spike-S1 and HKU1-spike-S1.**

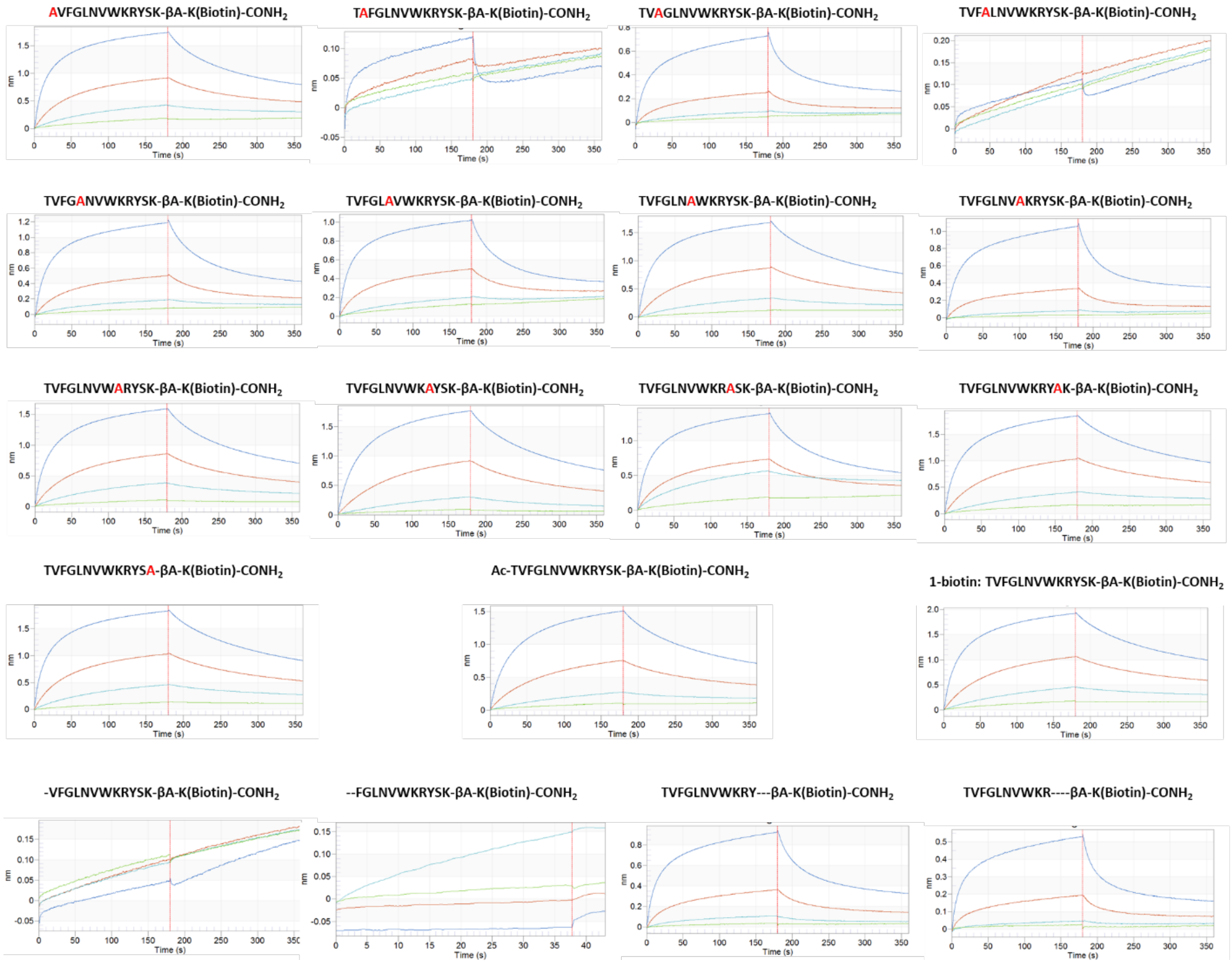

**SI Figure 8. BLI curves for alanine scan and truncation analysis of 1-biotin.** Association/dissociation of each peptide to SARS-CoV-2-spike-RBD was determined at four protein concentrations: 5000 nM, 1000 nM, 200 nM and 80 nM.

| | Sequence | Steady state Kd | Kinetic Kd | KD Error | kon(1/Ms) | kon Error | kdis(1/s) | kdis Error |
|---|---|---|---|---|---|---|---|---|
| 1 | TVFGLNVWKRYSK-X | 250 nM | 3.39E-07 | 3.38E-09 | 1.02E+04 | 8.46E+01 | 3.45E-03 | 1.91E-05 |
| 2 | LVMGLNAWNMWYK-X | 290 nM | 2.78E-07 | 2.87E-09 | 1.21E+04 | 1.04E+02 | 3.37E-03 | 1.94E-05 |
| 3 | LVOGLHVYLRQGK-X | 970 nM | | | | | | |
| 4 | LVMGLNVWLRYSK-X | 80 nM | 1.40E-07 | 2.73E-09 | 7.09E+03 | 6.23E+01 | 9.90E-04 | 1.73E-05 |
| | AVFGLNVWKRYSK-X | 270 nM | 5.16E-07 | 9.26E-09 | 8.66E+03 | 1.37E+02 | 4.47E-03 | 3.75E-05 |
| | TAFGLNVWKRYSK-X | No binding | | | | | | |
| | TVAGLNVWKRYSK-X | 1900 nM | 9.70E-07 | 2.88E-08 | 6.95E+03 | 1.89E+02 | 6.75E-03 | 8.08E-05 |
| | TVFALNVWKRYSK-X | No binding | | | | | | |
| | TVFGANVWKRYSK-X | 1200 nM | 8.91E-07 | 1.89E-08 | 7.38E+03 | 1.43E+02 | 6.57E-03 | 5.57E-05 |
| | TVFGLAVWKRYSK-X | 420 nM | 5.51E-07 | 2.04E-08 | 1.08E+04 | 3.68E+02 | 5.96E-03 | 8.73E-05 |
| | TVFGLNAWKRYSK-X | 500 nM | 6.32E-07 | 8.76E-09 | 7.15E+03 | 8.70E+01 | 4.52E-03 | 2.99E-05 |
| | TVFGLNVAKRYSK-X | 3200 nM | 1.13E-06 | 3.33E-08 | 7.13E+03 | 1.95E+02 | 8.04E-03 | 9.02E-05 |
| | TVFGLNVWARYSK-X | 380 nM | 6.69E-07 | 9.79E-09 | 7.26E+03 | 9.49E+01 | 4.86E-03 | 3.22E-05 |
| | TVFGLNVWKAYSK-X | 667 nM | 1.01E-06 | 1.14E-08 | 5.06E+03 | 5.11E+01 | 5.13E-03 | 2.56E-05 |
| | TVFGLNVWKRASK-X | 520 nM | 5.52E-07 | 1.57E-08 | 9.15E+03 | 2.37E+02 | 5.05E-03 | 5.91E-05 |
| | TVFGLNVWKRYAK-X | 270 nM | 5.21E-07 | 7.38E-09 | 7.04E+03 | 8.46E+01 | 3.67E-03 | 2.75E-05 |
| | TVFGLNVWKRYSA-X | 300 nM | 4.82E-07 | 7.51E-09 | 8.42E+03 | 1.14E+02 | 4.06E-03 | 3.13E-05 |
| | -VFGLNVWKRYSK-X | No binding | | | | | | |
| | --FGLNVWKRYSK-X | No binding | | | | | | |
| | TVFGLNVWKRY---X | 1900 nM | 8.67E-07 | 2.12E-08 | 8.22E+03 | 1.85E+02 | 7.13E-03 | 6.81E-05 |
|  | TVFGLNVWKR----X | 2500 nM | 1.10E-06 | 3.63E-08 | 8.16E+03 | 2.52E+02 | 8.96E-03 | 1.07E-04 |
| | Ac-TVFGLNVWKRYSK-X | 520 nM | 8.15E-07 | 1.10E-08 | 5.37E+03 | 6.35E+01 | 4.38E-03 | 2.89E-05 |
| | X = -bAla-Lys(Biotin)-CONH2 | | | | | | | |

**SI Table 1**. BLI affinity measurement result table for all peptides reported.

### Spike RBD pulldown from human serum

Bead preparation: Streptavidin coated magnetic beads (MyOne, 200 µL, 10 mg/mL) were washed with blocking buffer (PBS 1x = 0.05% tween-20, pH = 7.2, 3 x 1 mL) and then suspended in 1 mL of the same blocking buffer. **1-biotin** (20 µL, 0.5 mM) was added to the beads. After 30 minutes incubation at 4 °C the supernatant was removed and the beads washed with PBS (3 x 1 mL) and then resuspended in PBS (1 mL).

Human serum (100 µL, 7% protein content) was diluted with PBS (1x, pH = 7.2) to a final volume of 1 mL. CoV-2-spike-RBD (120 µL, 0.38 mg/mL) was mixed with diluted human serum (48 µL). Part of this mixture (80 µL) was added to the magnetic beads displaying immobilized **1-biotin**. This mixture was shaken for 45 minutes at 4 °C. The supernatant was removed and the beads washed with PBS at room temperature (5 x 1 mL, each wash 1 minute). The captured protein was eluted with 6 M urea solution (elution-1: 50 µL, 30 seconds and elution-2: 50 µL, 120 seconds).

For SDS-PAGE analysis 30 µL of each elution solution sample was loaded into the gel. In addition to the controls: CoV-2-spike-RBD and Human serum/CoV-2-spike-RBD. The analysis was performed using BoltTM 4-12% Bis-Tris Plus Gels (10-wells), 165 V for 36 min, utilizing pre-stained Invitrogen SeeBlueTM Plus2 molecular weight standard with BoltTM LDS Sample Buffer (4X).

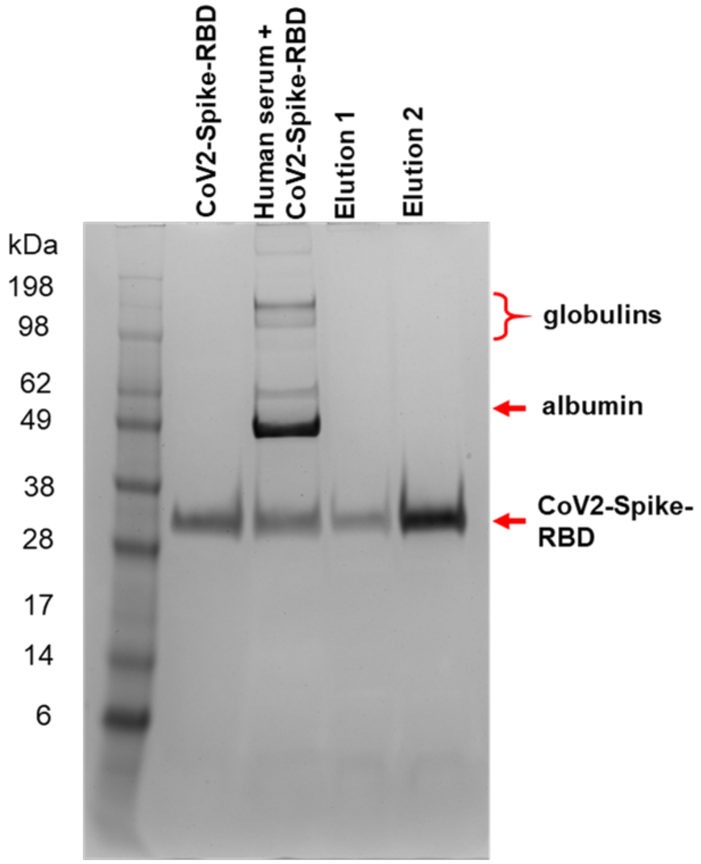

**SI Figure 9. SDS-PAGE analysis of the SARS-CoV-2-spike-RBD pulldown from human serum.**

### ELISA

**Buffer A (1x PBS, pH = 7.2 + 0.05% Tween 20)**

**Buffer B (10% FBS, 1% BSA, 0.05% Tween 20, 1x PBS, pH = 7.2)**

An ELISA plate (96-well format) was washed with buffer A (3 x 200 µL per well). SARS-CoV-2-Spike-RBD dissolved in buffer B in different concentrations was added (each concentration to three wells, each 100 µL) and incubated for 1 h at 37 °C. After this time, the supernatant was removed and the wells treated with blocking buffer (buffer B, 300 µL per well) and incubated overnight at 4 °C. Peptide 1-biotin (100 nM in buffer B) was added to all wells (beside the 'background' control wells) and incubated for 1 h at room temperature. The supernatant was removed, and the wells washed (3 x 300 µL buffer A per well). SA-HRP (0.1 µg/mL, 100 µL) was added to each well and incubated 30 minutes. All wells were washed (3 x 300 µL buffer A per well). TMB ELISA substrate was added to the wells (50 µL) and after 15 minutes quenched with sulfuric acid (50 µL). The absorbance at 450 nm was determined for all treated wells.

To detect RBD from human serum using our 1-biotin peptide, Sino Biological SARS-CoV-2-spike-RBD expressed from HEK293 cells was premixed with human serum at different concentrations (refer to figure 10), and then coated onto the ELISA plate overnight at 4 °C. Next day, the ELISA plate was blocked with 5% BSA in PBST (1 x PBS with 0.1% Tween-20) for 2.0 h at RT and then 1.0 h at 37 °C; Afterwards, the plate was treated with 1-biotin peptide at 100 nM in PBST, RT for 1.0 h, which then was followed by incubation with Streptavidin-HRP (0.1 µg/mL), RT for 30 mins. The signal was developed with ultra-TMB and then quenched with 2.0 M $H_2SO_4$, and finally, the absorbance at 450 nm was collected in a plate-reader. Between all steps, three washes were given with PBST (1 x PBS with 0.1 % Tween-20) with 5 min each wash.

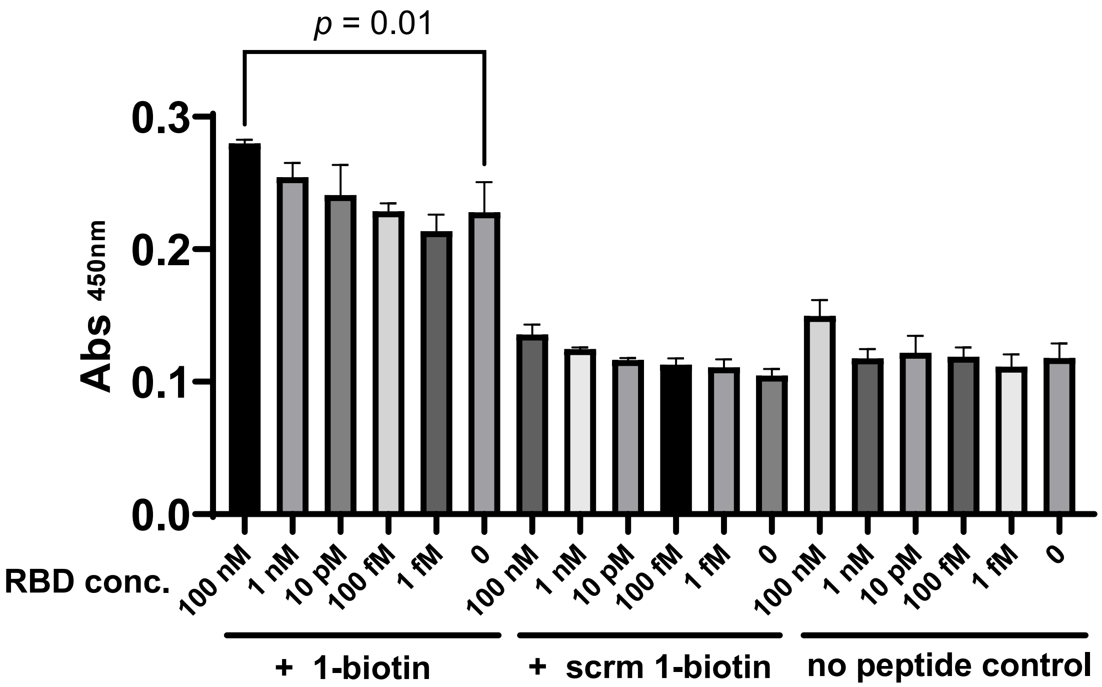

**Figure 10**. One hundred nanomolar of SARS-COV-2 Spike-RBD was detected from human serum. ELISA absorbance was taken at 450 nanometer. Measurements were performed in three technical repeats, and the significance was calculated with the unpaired t-test. RBD 100 nM vs no RBD: $p = 0.01$.
